# Sex-specific interactions between procedural and deliberative decision-making systems in a mouse model of Alzheimer’s disease

**DOI:** 10.1101/2021.03.16.435712

**Authors:** Rachel Anderson, Damyan W. Hart, Brian Sweis, Mathew A. Sherman, Mark J. Thomas, A. David Redish, Sylvain E. Lesné

## Abstract

A central question in aging and Alzheimer’s disease (AD) is when and how neural substrates underlying decision-making are altered. Here we show that while APP mice, a commonly used mouse model of AD, were able to learn Restaurant Row, a complex neuroeconomic decision-making task, they were significantly impaired in procedural, habit-forming, aspects of cognition and relied heavily on deliberation when making decisions. Surprisingly, these behavioral changes are associated with amyloid-beta (Aβ) pathology and network remodeling in the striatum, a key brain region involved in procedural cognition. Furthermore, APP mice and control mice relied on distinct sex-specific strategies in this neuroeconomic task. These findings provide foundational pillars to examine how aging and age-related neurodegenerative diseases impact decision-making across sexes. They also highlight the need for complex behavioral tasks that allow for the dissociation of competing neurally-distinct decision-making circuits to get an accurate picture of changes in neurodegenerative models of human disease.

## INTRODUCTION

Alzheimer’s Disease (AD) is a neurodegenerative disease clinically characterized by a progressive decline in cognitive skills, such as visual-spatial skills and memory, leading to dementia^1^. Although, historically, assessment of cognitive deficits in AD has focused on memory, language, and visuospatial skills^2^, growing attention is now being paid to how cognitive decision-making changes over the course of AD^3–5^. Elderly individuals have more difficulties than young individuals choosing between uncertain alternatives^6^ and more difficulties learning advantageous decision strategies^7^. These difficulties may be more pronounced when cognitive function is compromised by mild cognitive impairment (MCI) or by dementia^5, 8, 9^. While decision-making has been assessed in patients affected by neurodegenerative disorders such as Parkinson’s disease^9, 10^ or frontotemporal dementia^9, 11^, little is known about Alzheimer’s disease (AD) and its prodromal stage MCI beyond the fact that both are associated with gross impairments in decision-making^5, 9, 12^. In these human studies, assessment of decision-making has relied on paradigms such as the Iowa Gambling Task (IGT), where participants are tasked with maximizing bets under changing circumstances or the Cambridge Gambling Task (CGT), which assesses decision-making under risk. While informative, most of these studies have relied on simple assessments of risk or ambiguity, which do not capture other important aspects of decision-making such as value-based learning, foraging abilities, or deliberation. Importantly, these tasks capture fundamentally human measures of risky decision-making, which makes them difficult to assess in non-human animal models. In addition, these paradigms do not take into account modern theories that behavior arises from multiple decision-making systems^13–16^. Extensive work has shown that what appears to be similar behaviors on simple tasks could be produced by neurally-distinct computations^15, 16^. However, it is possible to identify behavioral markers within more complex tasks that can identify these neurally-distinct computations^16–18^. These different decision-making systems are instantiated in different neural circuits, and thus may become dysfunctional at different times in the development of disease. It is thus essential to identify the behavioral and economic phenotypes that account for individual variation in decision-making and to characterize the cognitive and motivational variables of intact and impaired decision-making. Further, advancing our basic knowledge of the cognitive and neuronal circuits underlying decision-making will also help us identify processes impacted in aging and in AD and other dementias.

Neuropathologically, AD is characterized by amyloid plaques composed of the amyloid-beta peptide (Aβ), neurofibrillary tangles (NFT) comprised of hyperphosphorylated tau proteins, neuronal death and neuroinflammation^19^. Brain Aβ deposition follows a pathological progression established by the Thal and Braak postmortem staging for regional extent of Aβ pathology^20^. Of note, these neuropathological studies as well as positron emission tomography (PET) amyloid imaging suggest that the presence of amyloid burden in the cerebral cortex alone is not an accurate predictor of clinicopathological AD^21^. Several groups have instead reported that striatal Aβ plaque density predicts the presence of a higher Braak NFT stage and clinicopathological AD in living subjects^22, 23^. Specifically, Hanseeuw and colleagues (2018) demonstrated that a novel three-category PET Aβ staging system that includes striatum better predicted hippocampal volumes and subsequent cognitive decline than a similar staging system including only cortical amyloid. Despite these exciting findings and the existence of previous work in individuals carrying familial AD mutations suggesting a strong relation between striatal Aβ and executive functions^24^, none of these studies determined whether Aβ pathology was linked to striatal dysfunction or whether some specific cognitive domains would be impaired. Additional work is thus needed to distinguish the respective contributions of striatal Aβ pathology to the onset of neuropsychiatric deficits or decision-making.

A number of mouse models have been developed that recapitulate some of these features of AD^25, 26^. Amongst those, the transgenic J20 mouse model^27^ overexpressing mutant human amyloid precursor protein (APP) has proven particularly useful when examining the role of Aβ on synaptic and memory deficits^28–32^ due to early plaque development, predominantly in the hippocampus at around 5-6 months of age^27, 33^. Recent work further expanded the characterization of amyloid pathology development in APP mice using whole brain imaging^33^, making this APP transgenic line an ideal candidate to study the impact of Aβ pathology on neural domains. Despite this seemingly extensive characterization of its phenotype, it is important to point out that only amygdala- and hippocampus-dependent memory modalities have been tested in this animal model, leaving unanswered questions of whether these mice are impaired in other decision-making modalities that either involve other brain structures or these same brain structures but engaged in multiple complex, dynamical ways.

Current theories of decision-making suggest that decisions arise from computationally separable systems implemented neurally through different neural circuits^14–16, 34^. Deliberative strategies depend on the ability to predict the consequences of one’s actions^17^. Spatially, these strategies depend on the presence of a cognitive map in the hippocampus containing information about the shape of the environment and the locations therein^15, 35^, however, evidence is that the cognitive “map” in the hippocampus is more general, containing general information about the structure of the world with which to plan^36–38^. In contrast, procedural strategies depend on well-practiced action-chains and fast pattern recognition of situations with stored cache values involving the dorsolateral striatum and the basal ganglia^16, 39^. Instinctual (Pavlovian) systems learn when to release actions from a limited action repertoire^16, 40^, involving the amygdala, periaqueductal gray, and nucleus accumbens shell^40, 41^. It is likely that the brain has evolved to have these different decision-making systems of which some are more advantageous in specific situations than others.

To better understand complex decision-making, we tested APP and control non-transgenic (nTG) littermates on a neuroeconomic spatial foraging task called Restaurant Row (RRow) that accesses multiple decision-making systems in controlled ways both within trial and across days. This neuroeconomic decision-making task was initially developed for rats^42^ and was recently adapted to mice^43^ and humans^44^ giving it immense translational value. Importantly, the RRow task^43^ allows for the dissection of different aspects of decision-making, including instinctual approach (Pavlovian), procedural habit (cached-action sequences), and deliberative (planning). While deliberative decision-making is thought to be hippocampal dependent, procedural decision-making relies on behavioral repetition and proper dorsolateral striatal functioning, and Pavlovian action-selection depends on amygdala function^15, 16, 40, 45^. The discrimination of these different sub-modalities of decision-making is particularly informative because when one of these neural circuits is impaired, another may compensate for it. For instance, as a consequence of age-dependent changes in hippocampal function, aged rats and humans shift from using a hippocampal-dependent “place” strategy to instead using a striatal-dependent “response” strategy when navigating^46–49^. Without knowing the strategy an animal is using, the gross behavioral output may look unimpaired while a disruption at the circuit-level may be overlooked^50, 51^. On the other hand, two similarly appearing behavioral impairments could arise from distinct circuit disruptions. Thus, pitting multiple decision systems against one another on a neuroeconomic task can read out competing processes that ultimately produce behavioral outputs and in turn aid in revealing the source of underlying computational dysfunction. Considering the well-documented impairment of hippocampal function in APP mice^29–32, 52–54^, we hypothesized that the multifaceted components of decision-making captured by RRow would be able to discern a more fine-grained approach to better characterize putative compensatory shifts in behavior.

In the present study, we examined young adult nTG and APP male and female mice on RRow at 6 months of age, when APP mice display intact spatial memory learning with impaired memory retention in hippocampal-dependent tasks such as the Barnes maze^53, 54^. All mice were able to learn this complex neuroeconomic decision-making task in which costs (delay to wait for food) start out low and then transition over weeks to become much higher (subsequently the reward environment becomes scarcer). Surprisingly, APP mice adapted their behavior more quickly upon the transition to scarcity and were able to renormalize their earnings to pre-transition numbers faster than nTG littermates. nTG mice typically accepted offers that, from prior experience with the different flavors, they most preferred and that were high in value (cheaper than what they were willing to spend in time waiting) quickly. This decision-making behavior resulted in less time deliberating, as has been observed previously^43^. By contrast, APP mice took significantly more time to decide before making a decision for all offers, whether advantageously cheap or disadvantageously expensive, and in general only took offers that they were willing to wait and earn based on their individual thresholds. This increased deliberative behavior was surprising considering that this APP transgenic mouse line is widely known for its hippocampal dysfunction. However, upon examination of Aβ plaque distribution throughout the brain, APP animals displayed a historically undocumented presence of amyloid deposits in the striatum. This new finding suggests that APP mice might have disrupted dorsolateral striatal procedural circuits. Overall, our results suggest that though APP mice were able to learn the RRow task, and perform it successfully, they required extensive deliberation to make any decision, even under conditions where their nTG counterparts used habitual, procedural decisions, quickly taking offers and then re-evaluating if necessary. Our results are the first to examine decision-making deficits across multiple decision-systems in a mouse model of AD and strongly emphasize the importance of examining multiple decision-making systems using tasks that access these multiple decision systems for behavior.

## RESULTS

### Hyperactivity and early discrimination of offers by APP mice

To better understand the cognitive and motivational behaviors of normative and impaired decision-making in AD, APP transgenic J20 mice and nTG littermates were subjected to RRow, a neuroeconomic decision-making task, to work for their sole source of food for the day (Fig. 1A). Mice were given one hour to traverse a square maze with four different feeding sites (i.e., restaurants), each with unique spatial cues and flavors. Upon entry into the “offer zone” (OZ, Fig. 1B) a tone indicated the delay animals would have to wait before getting a pellet. Higher pitch indicated longer delays while lower pitch indicated shorter delays. Delays were random on each entry, selected from a range depending on task stage (see methods). At this point, mice could choose to enter the “wait zone” (WZ) or skip the offer, thereby leaving the restaurant and continuing foraging by moving on to the next restaurant in the correct order (Fig. 1B). If the mouse decided to enter the WZ, the tone would step down in pitch until completion of the time delay, after which food would be delivered (earned). However, mice could re-evaluate their initial decision upon entering and quit the WZ at any time, forfeiting the pellet. Thus, the task was self-paced, and mice needed to alter their behavior to gain the most food in the one-hour time limit.

**Figure 1.**
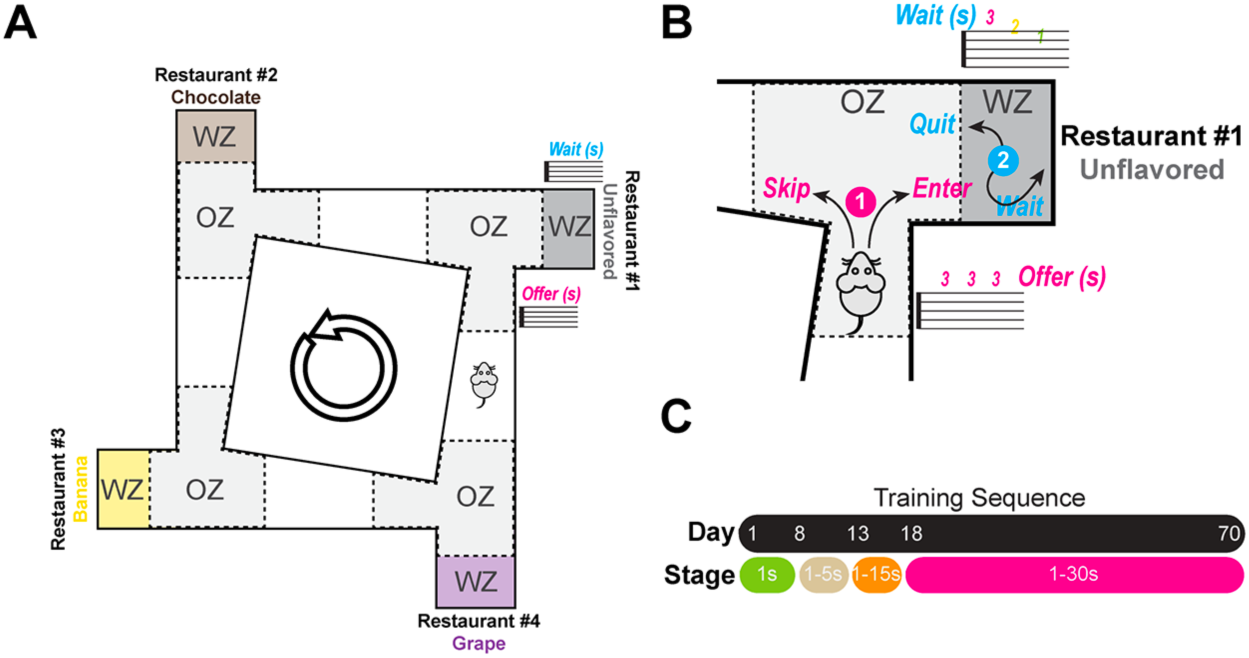
Restaurant Row Task. (**A**)Task schematic. Food-restricted mice (90% free food body weight) were trained to run counter-clockwise for flavored rewards in 4 “restaurants”. Restaurant flavor and location were identifiable with specific contextual cues and did not move location throughout the experiment. Each restaurant contained a separate offer zone (OZ) and wait zone (WZ). Tones sounded once the animal entered the offer zone. Fixed tone pitch indicated delay that mice would have to wait in the wait zone before earning a pellet. Upon entering the wait zone, tone pitch descended during delay “countdown”. Mice could quit the wait zone for the next restaurant during the countdown, terminating the trial. Mice were tested daily for 60 min in which they received their full food ration for the day. (**B**) Schematic of the offer (OZ) and wait zone (WZ). Initially in the OZ, mice have their first decision (1) where they can choose to accept the offer and enter the WZ or skip the offer and move on to the next restaurant. After entering the WZ, mice have another decision (2) where they can wait and earn the reward (flavored food pellet) or quit the WZ, not earning a pellet, and move on to the next restaurant. (**C**) Experimental timeline. Mice were trained for 70 consecutive days. Stages of training were broken up into blocks in which the range of possible offers began in a reward-rich environment (all offers were always 1 s, green epoch) and escalated to increasingly reward-scarce environments (offer ranges of 1–5 s, 1–15 s, 1–30 s).

Mice progressed from a reward-rich environment to a reward-scarce environment in stages across days (Fig. 1C). Each stage was defined by the range of possible delays that could be encountered upon entry into the OZ. The first stage spanned 7 days, during which all offers were only 1 s. Following the first stage, the range of offers encountered increased to 1-5 s. This second stage lasted 5 days (days 8-12), after which the offers increased to 1-15 s for the subsequent 5 days (days 13-17). The last stage (stage 4) spanned days 18-70 (i.e., rest of the experiment) and consisted of offers being randomly chosen between 1-30 s. Mice only had 1 hour to get all of their food for the day, so these changes in offer distributions produced increasingly reward-scarce environments.

This task was used previously with young 3-month-old C57BL/6J male mice to examine how non-transgenic mice behaved in a complex, neuroeconomic, decision-making task^43, 44, 55^. Here, we examined both male and female adult 6-month-old APP and C57BL/6J nTG littermates to determine the effect of early amyloid deposition in absence of overt neuronal or synaptic loss^27, 33, 53, 54, 56^ on decision-making neural substrates. Importantly at this age, APP mice display normal spatial reference learning but are slightly impaired spatial memory retention using the Barnes circular maze compared to control littermates^54, 57^.

All mice, including the seven APP mice and the eight nTG littermates, learned to run laps in the correct counterclockwise direction quickly during the first stage of the task (days 1-7, Fig. 2A). On day 1 and throughout the first week, APP mice ran significantly more laps than control mice (RM-ANOVA, *F*_(1,13)_ = 11.105, *p* = 0.0054; Fig. 2A and Suppl. Figure 1). As APP mice are well-known to display hyperactivity phenotypes^54, 56^, we measured running speed and distance travelled during the task. APP mice did travel more (Suppl. Figure 2) and ran faster during the first week of the experiment (Suppl. Figure 3), suggesting hyperactivity. This increase in average travel speed and distance equalized by the end of the experiment as nTG mice increased their running speed to match those of APP mice (Suppl. Figures 2A-3A) suggesting that by the later stages of the experiment, differences in reward-rate were not due to running speeds. During the first week (1 s offers), APP mice also earned significantly more pellets than controls (RM-ANOVA, *F*_(1,13)_ = 9.777, *p* = 0.008; Fig. 2B and Suppl. Figure 4). However, the two groups showed similar earning rates in the subsequent 2^nd^ and 3^rd^ stages of the task (see below), even though APP mice were still covering more ground and running faster than nTG mice (Suppl. Figures 2A-3A). All mice developed flavor preferences in the first stage of the experiment, whose rank order remained stable throughout the experiment (Suppl. Figure 5).

**Figure 2.**
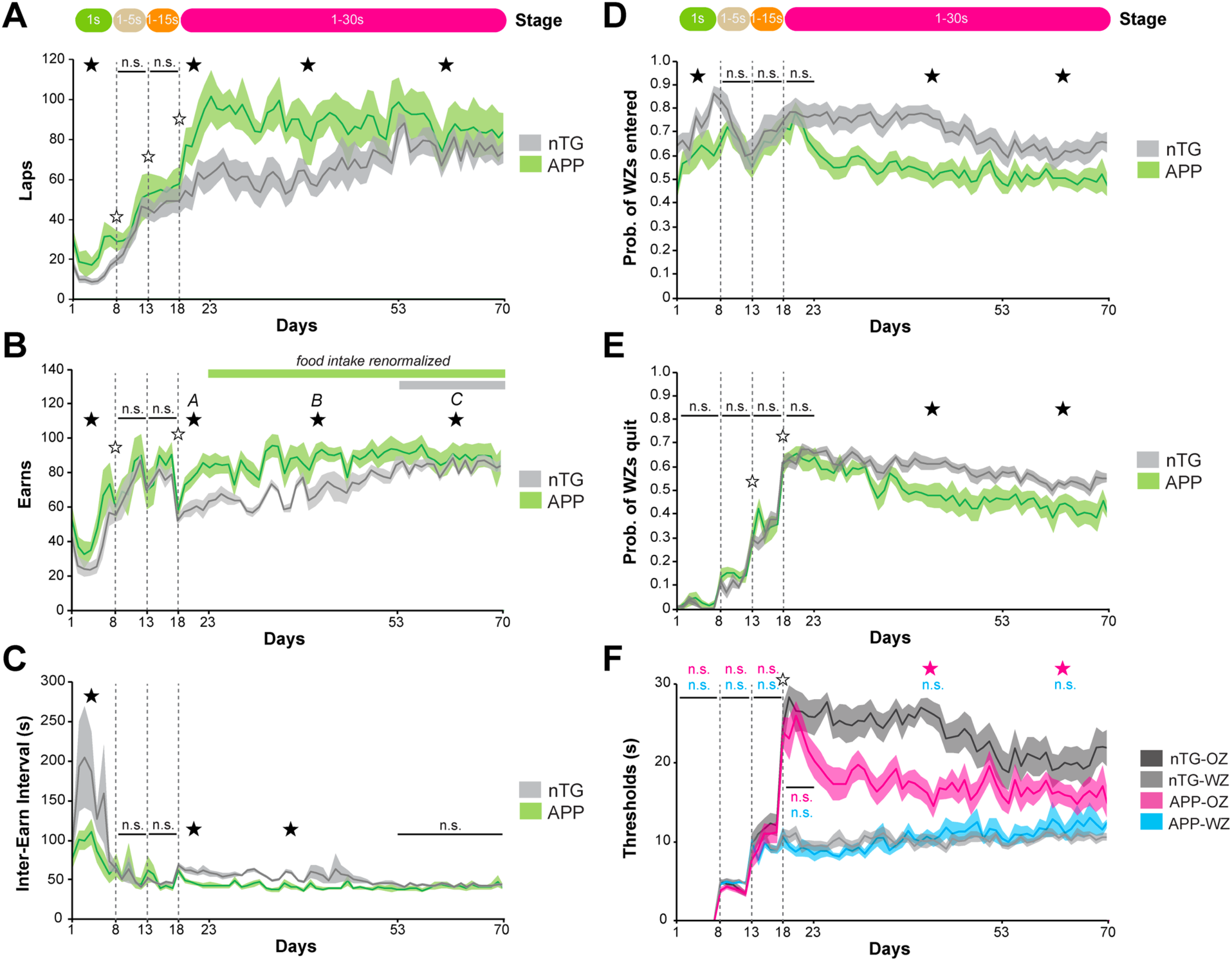
APP mice show distinct behavioral differences compared to nTG mice. (**A**) Number of correct (counterclockwise) laps run. (**B**) Total number of earns. Transition to the 1-30 s offer block resulted in a significant decrease in earns for both groups. By day 23, APP mice renormalized their earnings back to levels compared to previous stages, whereas nTG mice did not restabilize their earnings until day 53. (**C**) Reinforcement rate measured as average time between earnings. (**D**) Proportion of total wait zones (WZs) entered versus skipped. (**E**) Proportion of total WZs earned versus quit. (**F**) Offer zone (OZ) decision thresholds and wait zone (WZ) decision thresholds as a function of cost (offer delay). Early in training, OZ and WZ thresholds are equivalent, indicating that mice entered offers that they were in turn willing to wait for. Following the transition to 1-30 s offers, OZ thresholds were much higher than WZ thresholds, indicating that mice entered offers that they were not willing to wait and earn. Data are presented as the daily means (± SEM) across the entire experiment. The x-axis reflects the days of training with vertical lines indicating changes in training stages (corresponding with Fig. 1B). White start on a vertical transition line indicates immediate significant behavioral change at the block transition within groups; black star indicates significant differences between groups within the given training stage.

**Figure 3.**
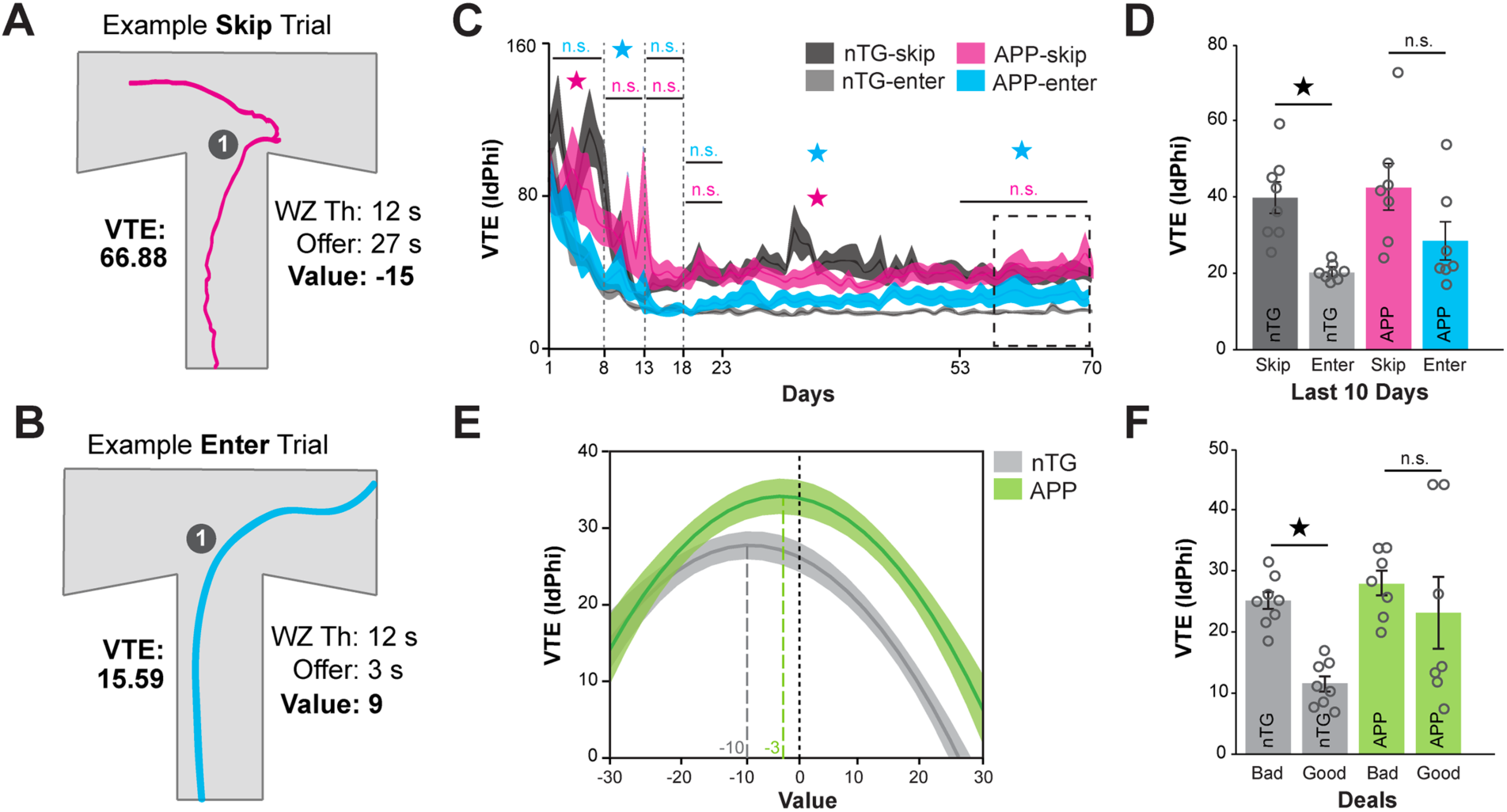
Vicarious-Trial-and-Error. (**A**) Example of a nTG mouse’s path trajectory in the offer zone (OZ, decision point 1) during a single skip trial (from day 70). Mice initially orient toward entering (to the right) but then ultimately reorient and skip. This orientation is the basis for the IdPhi measurement used to classify vicarious-trial-and-error (VTE). Here we show how value of the offer is calculated based on the offer (in this case 27 seconds) and the individual’s willingness to wait (WZ threshold, in this case 12 s). Value is calculated by subtracting the WZ threshold from the offer (12-27 = −15). (**B**) Example of a nTG mouse’s path trajectory in the OZ (decision point 1) during a single enter trial (from day 70). nTG mice make relatively smooth entrances into the WZ while APP mice show sharper trajectories leading to higher VTE values when entering. Here again we illustrate how the value is calculated (WZ threshold - offer; in this case 12-3 = value of 9). (**C**) Average VTE (IdPhi) values split by skip versus enter decisions across days of training by groups (dark grey, grey = nTG, pink and blue = APP). Pink star denotes p<0.05 for between-group differences in skip VTE. Blue star denotes p <0.05 for between group differences in enter VTE. Data are presented as the daily means (± SEM) across the entire experiment. Vertical dashed lines represent stage transitions. (**D**) Bar graph depicting mean ± SEM for well-trained animals (last 10 days) to illustrate group differences. *, p<0.05 one-way ANOVA within group (nTG). (**E**) Quadratic functions of VTE by value (WZ Th. minus offer) for the last 10 days. Negative value denotes an economically unfavorable offer. Peaks are right-shifted for APP mice (value of −3 compared to −10 for nTG mice), indicating higher VTE for closer to neutral values. (**F**) Mean ± SEM of values from E for bad (value <0) and good (value >0) deals *, p<0.05 one-way ANOVA within group (nTG).

To determine how efficiently mice were using their time, we analyzed reinforcement rate as the amount of time in seconds between earning a pellet (inter-earn-interval, IEI). During the first week of the task, APP mice had a shorter IEI compared to nTG animals (peaking at ∼140s vs. ∼250s respectively), indicating higher reinforcement rates in APP mice than in control littermate mice (RM-ANOVA, *F*_(1,13)_ = 10.35, *p* = 0.0018; Fig. 2C), though this equalized between groups in the next 10 days of the task.

The second stage of the experiment (beginning day 8) consisted of offers varying between 1-5 s and lasted for 5 days (days 8-12). The number of laps run and pellets earned increased from the previous stage (RM-ANOVA between stages indicated by the open star on dotted lines; laps: *F*_(1,13)_ = 42.93, *p* < 0.0001; earns: *F*_(1,13)_ = 12.692, *p* = 0.0035; Fig. 2A, B) but equalized between the groups, and subsequently the reinforcement rates stabilized and equalized between APP and nTG mice (Fig. 2C). Following 1-5 s offers, mice transitioned to the third stage consisting of 1-15 s offers between days 13-17. All mice continued increasing the number of laps they ran, and both groups did this equally (RM-ANOVA between stages indicated by the open star on dotted lines; *F*_(1,13)_ = 25.22, *p* = 0.0002; Fig. 2A). Though mice were running more laps, the number of pellets earned remained stable relative to the previous stage (Fig. 2B).

A key element to RRow is that it entails multiple junctures of decision-making, allowing us to examine different behavioral components involved in these decisions (Fig. 1B). Upon entering the OZ, mice can choose to accept the offer and enter the WZ or decide to skip and continue on to the next restaurant. Mice who enter the WZ can then decide to wait out the delay thus earning food or can re-evaluate and quit, forfeiting the pellet and continuing on to the next restaurant (Fig. 1B). Prior work^43^ has suggested that the decision to skip or enter the WZ develops differently than the decision of whether to wait and earn or quit out of the WZ. Non-transgenic mice entered WZs for most offers that were presented to them, regardless of flavor, during the first week of the experiment when offer durations were very short (Fig. 2D, *grey line*). This pattern of accepting most offers was starkly different from APP mice who began discriminating among the offers they accepted and instead starting skipping offers (RM-ANOVA, *F*_(1,13)_ = 7.801, *p* = 0.016; Fig. 2D). As costs began to increase with the progression of stages, nTG began entering WZs at the same rate as APP mice (Fig. 2D).

With the increase in offer length, all mice also began quitting accepted offers (RM-ANOVA between stages 2 and 3 indicated by the open star, *F*_(1,13)_ = 6.062, *p* = 0.029; Fig. 2E). Lastly, we analyzed the thresholds of the willingness to enter an offer (OZ threshold), as well as the willingness to wait for the reward (WZ threshold) to assess how mice handled offers below or above their individual threshold. During the 1-5 s (2^nd^) and 1-15 s (3^rd^) stages while the reward environments were still relatively rich, OZ and WZ thresholds were equivalent in both groups, indicating that all mice, for the most part, decided to enter wait zones where the delay matched how long they were willing to wait and earn (Fig. 2F).

Together, these data from the initial stages of the experiment are consistent with prior work relying on this task despite using older adult animals^43^. These results suggest that both APP and nTG littermates were able to learn this complex neuroeconomic decision-making task and to adapt to evolving scarcity.

### APP Mice Adapt to Scarce Foraging Environment Faster than Control Mice

Upon transitioning to the final stage in which offers ranged from 1-30 s, all mice experienced a drop in the number of pellets earned as they learned to navigate the increased offer lengths (RM-ANOVA between stages 3 and 4, *F*_(1,13)_ = 26,49, *p* = 0.0002, indicated by the open star, Fig. 2B). Almost immediately (on the second day of this stage, day 19 of the task), APP mice responded to this new environment, ran more laps, and were able to earn significantly more pellets than controls (RM-ANOVA, laps: *F*_(1,13)_ =4.022, *p* =0.06; earns: *F*_(1,13)_ = 8.018, *p* = 0.014, Fig. 2A, B). In fact, APP mice were able to renormalize their earnings in the reward-scarce environment to the earnings they had achieved in the reward-rich environments within a couple of days, i.e., by day 23 of the task (Fig. 2B, *green bar*). In contrast, nTG littermates did not reach pre-transition earnings until day 53 of the task (Fig. 2B, *grey bar*). Using this information, we analyzed this fourth and final stage of the experiment in three separate epochs. The first epoch consists of days 18-22 (epoch A). In this epoch both groups have yet to renormalize their earnings to previous stages. In the second epoch (days 23-52, epoch B) APP mice have sufficiently renormalized their earnings but nTG mice have yet to earn as much as they did in stage 3 of the task. In the final epoch (days 53-70, epoch C) all mice have renormalized their food intake.

Prior to this final stage (before day 18) where the reward-distribution became scarce, OZ and WZ thresholds for both groups were equivalent, as mice generally took offers for which they were willing to wait and earn. However, immediately following the transition to 1-30 second offers, all mice initially took most of the offers in the OZ as indicated by the very high 20-25 s OZ thresholds for both groups that increased on day 18 (RM-ANOVA for transition between stages 3 and 4, *F*_(1,13)_ = 67.56, *p* < 0.0001, indicated by the open star; Fig. 2F, *black and pink lines*) but quit the majority of trials that were accepted with a starting delay above 10 s as indicated by the ∼10 s WZ thresholds, suggesting offer zone decisions were not in register with willingness to wait in the wait zone (Fig. 2F, *grey and blue lines*). Over time and despite the scarcity of the environment, all mice developed lower OZ thresholds, indicating that the delay for which they were willing to enter the WZ approached the delay for which they were willing to wait in the WZ (Fig. 2F). Here again, APP mice adapted their behavior to this change in reward scarce environment quickly by skipping more offers than nTG mice by epoch B (days 23-52, RM-ANOVA, *F*_(1,13)_ = 288.42, *p* < 0.0001; Fig. 2D). APP mice continued skipping more offers than nTG for the rest of the experiment (Epoch C days 53-70, RM-ANOVA, *F*_(1,13)_ = 81.944, *p* < 0.0001; Fig. 2D). By not entering WZs for high offers, APP mice showed a quick drop in OZ thresholds that became significantly lower than the OZ thresholds of nTG mice during epoch B of the last stage of training (days 23-52, RM-ANOVA, *F*_(1,13)_ = 223.16, *p* < 0.0001; Fig 2F) and that eventually became in register with WZ thresholds (Fig. 2F). APP mice had significantly lower OZ thresholds than nTG mice during the rest of the experiment, even after nTG mice renormalized their food intake (epoch C, days 53-70, RM-ANOVA, *F*_(1,13)_ = 39.31, *p* < 0.0001; Fig. 2F).

The fact that by the end of the experiment, OZ and WZ thresholds were not significantly different in APP mice suggests that APP mice generally entered the WZ when the offer was one that they would wait to earn. This hypothesis was further supported by examining how often APP mice quit upon entering the WZ. Consistent with this observation, APP mice quit significantly less than control mice starting during epoch B (days 23-52, RM-ANOVA, *F*_(1,13)_ = 97.94, *p* < 0.0001; Fig. 2E) and continuing throughout the completion of the task (Epoch C, days 53-70, RM-ANOVA, *F*_(1,13)_ = 84.01, *p* < 0.0001; Fig. 2E).

Together, these data suggest that APP mice were more selective in the offers they chose to enter and were more likely to wait and earn pellets upon entering, whereas control mice were more likely to enter but also more likely to quit.

### Mice Show Vicarious-Trial and Error in the Offer Zone

To better understand how mice approached the decision to enter or skip offers in the task, we measured deliberative behavior. To assess ongoing deliberation and planning at the decision point (OZ), we examined vicarious-trial and error (VTE), a behavioral phenomenon in which an animal pauses at a choice point and orients sequentially towards its options. VTE was estimated by calculating the absolute integrated angular velocity, *IdPhi*, whereby larger *IdPhi* corresponds to increased VTE^58^. Prior studies using RRow found that mice showed increased VTE (more biphasic trajectories, which include a sharp turn) when making the decision to skip, with less VTE (smooth entrances) when eventually deciding to enter^43^ as can be seen in illustrations of examples from a nTG mouse from this study (Fig. 3A, B). When the environment was rich in rewards during stages 1-3, and as animals increased their knowledge of the task, VTE enter and VTE skip scores equally decreased from ∼80 to ∼20-40 for both nTG and APP groups (Fig. 3C). This is consistent with previous observations^43^.

Though previous observations suggest the decision to skip required more VTE than the decision to enter^43^, we did not see this in APP mice. APP mice presented with an offer they would go on to accept showed a biphasic trajectory similar to what we would expect of an eventual skip. By the epoch B (days 23-52) of the last stage of the task (1-30 s offers), APP mice showed higher VTE values when entering WZs than nTG mice (RM-ANOVA, *F*_(1,13)_ = 80.05, *p* < 0.0001; Fig. 3C). APP mice consistently deliberated more upon making a decision to enter than nTG throughout the rest of the experiment (epoch C, days 53-70, RM-ANOVA, *F*_(1,13)_ = 49.61, *p* < 0.0001, indicated by the blue star; Fig. 3C). In contrast, VTE skip values were largely equalized between groups, except for Epoch B (days 23-52) in which nTG mice showed greater VTE values when making the decision to skip (RM-ANOVA, *F*_(1,13)_ = 27.02, *p* < 0.0001, indicated by the pink star; Fig. 3C).

By the last 10 days of the experiment, when all animals had renormalized food and were making consistent behavior choices, nTG mice showed significantly lower VTE amounts when entering, than when deciding to skip (one-way ANOVA within-group *F*_(1,7)_ = 24.08, *p* = 0.0002; Fig. 3D). In contrast to the discrepancy nTG mice showed when skipping or entering, APP mice showed equal levels of VTE whether eventually entering or skipping (one-way ANOVA within-group *F*_(1,6)_ = 3.222, *p* = 0.098). Student t-tests within- and between-groups (nTG enter, nTG skip, APP enter, APP skip) revealed significant differences between nTG enter and nTG skip (F_(1,7)_ = 24.08, *p* =0.0002; Fig. 3D) but no other significant group differences. Together, the VTE results suggested that, for nTG mice, the decision to enter the WZ or the decision to skip the offer in the OZ was dichotomous but APP mice appeared to treat the two decisions equally, requiring high deliberation upon making any decision.

Because previous data have demonstrated that low-value offers (where the delay is expensive relative to the individual’s threshold) require more deliberation and more time to decide than high-value offers (where the delay is cheap relative to the individual’s threshold) across species^44^, we next evaluated the amount of VTE based on the value of the offer. We defined the value of the offer by subtracting the given offer from the WZ threshold for that animal, i.e., value= WZ Th – Offer (concrete examples are provided in Fig. 3A, B). Across the spectrum of possible values, VTE distributions for both groups followed a quadratic function (last 10 days of the experiment; Fig. 3E), consistent with previous studies^43^. However, APP mice showed larger *IdPhi* values at peak than nTG mice (Fig. 3E) indicating that they showed more VTE. Importantly, the value at which the peak of the *IdPhi* distribution appeared was shifted to the right (higher value, better offer) for APP mice relative to nTG mice (−3 value compared to nTG value of −10). Segregating value offers between economically good deals (Value > 0) and bad deals (Value < 0) for the last 10 days when animals were well trained, control animals displayed markedly higher VTEs for bad offers and lower VTE for good offers (One-way ANOVA within group *F*_(1,7)_ = 48.615, *p* < 0.001; Fig. 3F). This makes intuitive sense as it is a sound economic decision to take good, cheap offers. By contrast, turning down an offer, even if expensive, may take more processing when considering the other option is exploring other unknown opportunities. It may take more mental effort to leave a known, though expensive, reward in a scarce environment.

Importantly, APP mice did not show this distinction and instead showed similar amounts of VTE (One-way ANOVA, *F*_(1,6)_ = 0.601, *p* = 0.453; Fig. 3F) prior to deciding, regardless of whether the offer was good or bad. Analyses between groups for both good and bad values showed that while nTG and APP mice deliberated equally for bad offers (Fig. 3F), APP mice showed higher VTE (*F*_(1,14)_ = 4.338, *p* = 0.05; Fig. 3F) for good offers than nTG mice. These data further support the conclusion that nTG mice deliberated more when they had to make skip decisions in conflict with their approach desires, but that APP mice deliberated more regardless of whether the offer presented was economically cheap or expensive, or whether they skipped or entered.

This leads to the question as to why APP mice deliberated more for good offers? In a scarce reward environment, good offers (value > 0) should be taken. nTG mice reliably demonstrate this, with low VTE (smooth entrances) for values greater than 0. By the end of the experiment, this should be procedural and habitual. The fact that APP mice did not show smooth entrances for good offers, and instead showed high amounts of deliberation (equal to amounts of bad offers) led us to question if APP mice had impaired procedural decision-making. To investigate this, we examined the amount of VTE animals demonstrated for distinct flavor preferences. All animals showed reliable flavor preferences throughout the experiment, earning more of their preferred flavor than their least preferred flavor (Suppl. Figure 5; nTG: first: *F*_(3,220)_ = 6.32, *p* = 0.0004; second: *F*_(3,156)_ = 38.92, *p* < 0.0001; third: *F*_(3,156)_ = 98.49, *p* <0.0001; epoch A: *F*_(3,156)_ = 67.69, *p* <0.0001; epoch B: *F*_(3,956)_ = 355.9, *p* < 0.001; epoch C: *F*_(3,572)_ = 265.1, *p* < 0.001; APP: first: *F*_(3,192)_ = 20.12, *p* < 0.001; second: *F*_(3,136)_ = 33.75, *p* < 0.0001; third: *F*_(3,136)_ = 68.88, *p* < 0.0001; epoch A: *F*_(3,136)_ = 58.15, *p* <0.0001; epoch B: *F*_(3,836)_ = 342.2, *p* < 0.0001; epoch C: *F*_(3,500)_ = 209.5, *p* < 0.0001). Based on previous reports^43^, mice showed distinct differences in VTE for their most and least preferred flavors. Collapsed across days, nTG mice showed very little VTE for their most preferred flavor, indicating smooth entrances, and deliberated much more before deciding for their least preferred flavor (*F*_(3,28)_ = 6.47, *p* = 0.002). Surprisingly, APP mice did not show different levels of deliberation for different flavor preferences (*F*_(3,24)_ = 0.26, *p* = 0.85) and instead showed relatively consistent, high levels of VTE for all flavors.

When we evaluated VTE for flavor preferences by stages throughout the experiment, nTG mice showed differences in VTE by flavor for every stage, except the first stage where costs were low (Suppl. Figure 6A; first stage, *F*_(3,196)_ = 1.55, *p* = 0.2; second stage, *F*_(3,140)_ = 7.68, *p* < 0.0001; third stage: *F*_(3,140)_ = 14.17, *p* < 0.0001; epoch A: *F*_(3,140)_ = 4.45, *p* = 0.005; epoch B: *F*_(3,840)_ = 26.01, *p* < 0.0001; epoch C: *F*_(3,504)_ = 8.53, *p* < 0.0001). When analyzed across the experiment, we observed that APP mice demonstrated distinct VTE levels for different flavors, showing higher deliberation for their least preferred flavor and lower VTE for their most preferred flavor, similar to nTG mice, for the first part of the experiment (Suppl. Figure 6B, first stage, *F*_(3,164)_ = 0.74, *p* = 0.53; second stage, *F*_(3,120)_ = 7.39, *p* =0.0001; third stage: *F*_(3,120)_ = 6.62, *p* = 0.0004; epoch A: *F*_(3,120)_ = 5.13, *p* = 0.002). However, by the time they renormalized their food intake (day 23), APP mice stopped showing this distinction and instead deliberated equally for all flavors, disregarding flavor preference, though still earning more of their preferred flavor (epoch B: *F*_(3,720)_ = 1.2, *p* = 0.31; epoch C: *F*_(3,432)_ = 1.2, *p* = 0.31).

### Vicarious-Trial and Error Used Differently throughout the Experiment

Time spent in the OZ is wasted time from the time budget. It would be economically beneficial for mice to enter the WZ directly, make the decision while in the WZ, and then quit as soon as they realized they had entered a bad deal. However, previous work has found that neither mice, rats, nor humans behave this way^42, 43, 58^, and that instead all are sensitive to aspects of choice history beyond the value strictly tied to the reward itself. Additional information, which may be associated with fluctuations in affective state, is also taken into consideration in the OZ and WZ. Previous work has found that OZ time does reduce the efficiency of getting food on the RRow task^43^. However, we observed that APP mice displayed enhanced deliberative processes (i.e., higher VTE) and earned more food than nTG mice, suggesting that the APP behaviors produced an increased efficiency. To determine the role VTE was playing in both the nTG and APP mice, we simulated the number of earns one would expect if the mice had hypothetically displayed different VTE behaviors in relation to OZ choice outcome. First, we identified high-VTE vs. low-VTE trials for each mouse defined by a median split among all VTE values for each day, as previously reported^43^. We then took two approaches to simulate hypothetical alternatives to the total number of pellets earned: replace the trial outcome of high-VTE trials with that of low-VTE trials or simply force all high-VTE trials to end with no food earned before calculating expected earns for that day. Finally, we then looked at all of the stages of the experiment, including the three epochs in the last stage (epoch A, no food renormalization: days 18-22; epoch B, APP renormalized: days 23-52; epoch C, all animals renormalized: days 53-70) to see how patterns in VTE are related to the number of earns throughout the process of mastering the task (Fig. 4A, B).

**Figure 4.**
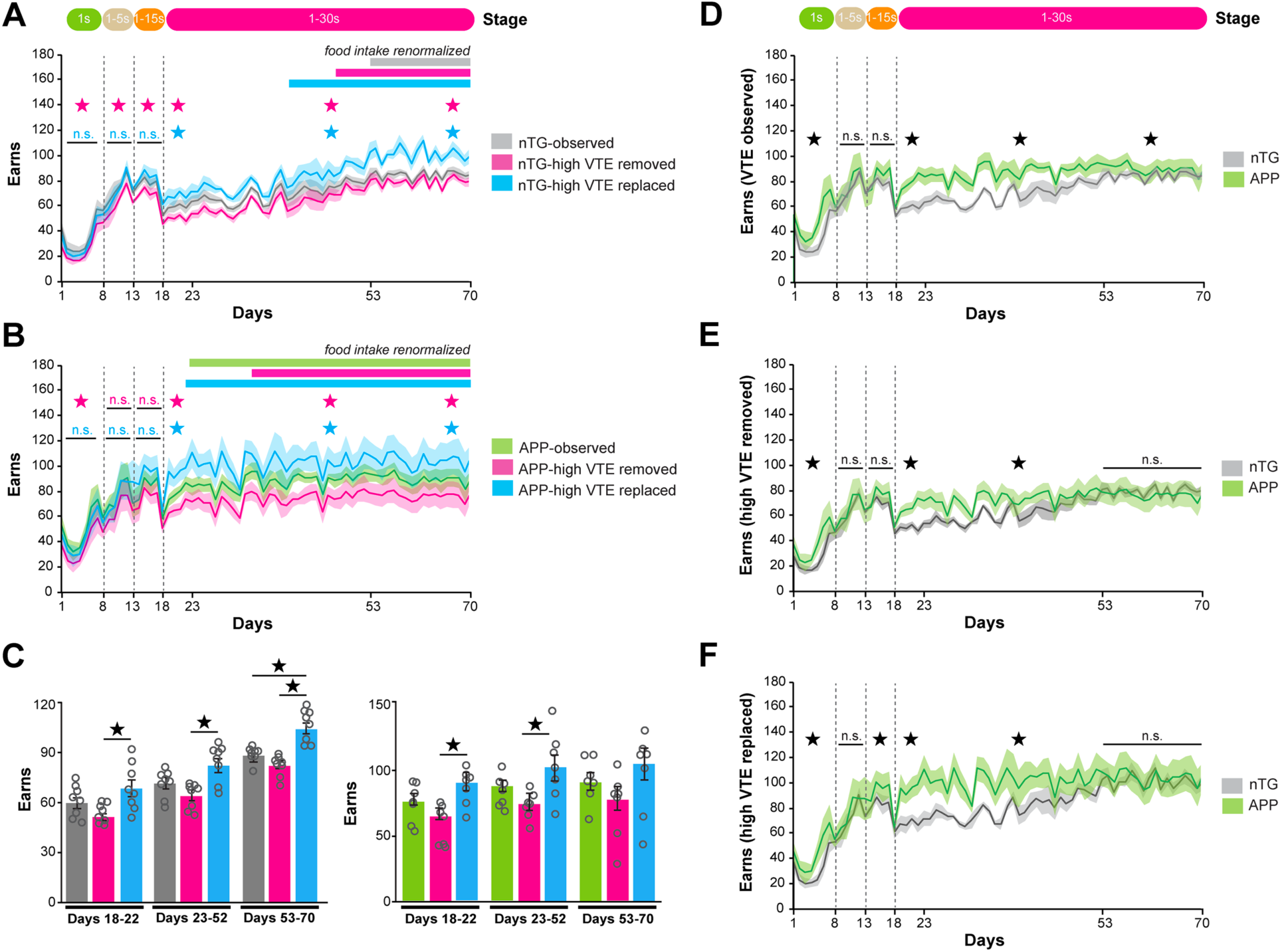
Earn simulations. (**A,B**) Earn simulations split by group (nTG, APP respectively). High VTE trials were determined as being greater than the median VTE for an individual mouse on that day. High VTE trials were first removed and earns were analyzed (removed; pink). Then high-VTE trials were replaced with low-VTE trials (VTE trials < median; replaced, light blue). nTG (*A*) and APP mice (*B*) showed an effect on earning potential based on simulation during the last stage of training, earning significantly more pellets if high VTE trials were replaced with low VTE trials (denoted by light blue star), and earning significantly less when high VTE trials were removed all together (denoted by pink star). Data are presented as the daily means (± SEM) across the entire experiment. Vertical dashed lines represent stage transitions. (**C**) To look at how earnings changed in these simulations based on training, we examined average earnings by simulation during three separate epochs of the last stage of training (when no animals had their food intake renormalized: days 18-22; when APP had their food intake renormalized: days 23-52; when all animals had their food intake renormalized: days 53 to 70). nTG mice had significantly higher earning potential when high VTE events were replaced with low VTE events in the last epoch (days 53-73; Fig. 4C left). In contrast, APP mice do not show any significant differences in earning simulations when the earnings are averaged across days (right). Data are presented as mean ± SEM. (**D**) Actual earns that were observed in this experiment. APP mice earned significantly more than nTG mice during the last stage of training (1-30 second offers; RM-ANOVA between groups). (**E**) High VTE removed simulations compared between groups. Simulated APP mice earn significantly more early on in the last stage of training but nTG mice catch up to APP mice during the last epoch (days 53-70). (**F**) High VTE replaced with low VTE simulations compared between groups. Simulated APP mice earn significantly more early in the last stage of training (1-30 s offers) but nTG mice catch up by the end (days 53-70). Data are presented as the daily means (± SEM) across the entire experiment. Vertical dashed lines represent stage transitions.

For nTG mice, replacing high VTE trials with low VTE trials increased their earnings during the last stage of training (RM-ANOVA epoch A: *F*_(1,13)_ = 8.113, *p* = 0.005; epoch B: *F*_(1,13)_ = 61.662, *p* < 0.0001; epoch C: *F*_(1,13)_ = 143.75, *p* < 0.0001; indicated by the blue star; Fig. 4A). Simulated nTG mice who had high VTE trials replaced with low VTE renormalized their food intake by day 39 (compared to the observed renormalization of day 53). However, in nTG mice, completely removing high VTE trials (all VTE > median split for that mouse) altogether resulted in a loss of earns compared to what we observed (RM-ANOVA epoch A: *F*_(1,13)_ = 13.13, *p* = 0.0005; epoch B: *F*_(1,13)_ = 39.21, *p* < 0.0001; epoch C: *F*_(1,13)_ = 19.14, *p* < 0.0001; indicated by the pink star; Fig. 4A). Interestingly, in simulated nTG mice, removing high VTE trials renormalized their food intake faster than the observed mice (day 48 compared to day 53) though the number of earns they needed to get back to was lower (70 daily earns compared to 80 daily earns).

For APP mice, the same trend replicated. Replacing high VTE trials with low VTE trials significantly increased their earnings during the last stage of training (RM-ANOVA epoch A: *F*_(1,13)_ = 11.89, *p* = 0.001; epoch B: *F*_(1,13)_ = 37.89, *p* < 0.0001; epoch C: *F*_(1,13)_ = 18.11, *p* < 0.0001; indicated by the blue star; Fig. 4B). Simulated APP mice with replaced VTE trials showed renormalized earnings just one day earlier than observed mice (day 21 versus 22). When high VTE trials were completely removed, APP mice were penalized with loss of earns (RM-ANOVA epoch A: *F*_(1,13)_ = 11.85, *p* = 0.0011; epoch B: *F*_(1,13)_ = 54.31, *p* < 0.0001; epoch C: *F*_(1,13)_ = 20.15, *p* < 0.0001; indicated by the pink star; Fig. 4B). Unlike nTG mice, who still exhibited renormalized food faster without high VTE trials at all, simulated APP mice took longer to renormalize food than the observed APP mice (renormalizing by day 30 instead of day 23), showing an extreme penalization.

ANOVA analyses comparing the means of earns during the last stages revealed that, for nTG mice, replacing high VTE had the most impact on earning potential (Fig. 4C), particularly during the last epoch (epoch C; *F*_(2,21)_ = 20.72, *p* < 0.0001) with simulated nTG animals who had high VTE trials replaced with low VTE trials earning more than observed mice (post-hoc analysis: *p* = 0.0003). APP mice did not show significant differences when the earnings are averaged and compared between simulations (Fig. 4C).

Lastly, when simulations were compared by condition (Fig. 4D-F), it became clear that nTG mice benefit from high VTE trials removed or replaced with low VTE trials. In the observed data, APP mice earn more than their nTG littermates throughout the last stage of the experiment (Fig. 4D). However, in both removed (Fig. 4E) and replaced simulations (Fig. 4F), nTG mice are able to earn as much as their APP counterparts by the last epoch (days 53-70). These data imply that some deliberation is important for maximum earnings but that too much deliberation is costly, for both nTG and APP mice. However, it is important to note that the impact of deliberation was stronger for nTG mice who had much more to gain by deliberating less than their median amount. This also suggests that VTE may be serving other purposes beyond just reinforcement maximization in nTG mice.

### Newly identified striatal Aβ deposition in APP mice

Because amyloid plaques are already present in the parenchyma of APP mice at 9 months of age^27, 31, 33, 56^ (age at which mice ended the RRow task), we wondered whether development of amyloid-beta (Aβ) pathology could explain the altered behavior of APP animals compared to nTG littermates. Despite the extensive use of J20 APP transgenic mice as a model of AD, the spatiotemporal distribution of amyloid pathology in this model has only been sporadically characterized (https://www.alzforum.org/research-models/j20-pdgf-appswind). While recent work provided a whole-brain assessment of methoxy-X04-positive plaques in this line^33^, the approach used inherently missed early amyloid pathology present at the onset of cognitive deficits in hippocampus-dependent tasks^54, 57^. To fill this knowledge gap and to establish the association of possible pathologically impacted neural networks with underlying behavioral changes in RRow, we sought to gain a better understanding of the overall localization of amyloid deposits in APP mice using an unbiased approach. Immediately following the RRow task, brains from 9-month-old APP and nTG mice were mass-processed using MultiBrain^®^ technology and analyzed by immunolabeling using the antibody 6E10 to spatially characterize amyloid plaque localization (Fig. 5A). No staining was observed when the antibody was used to stain tissue from nTG mice. Moreover, each Aβ plaque was confirmed morphologically by examination at high-power magnification (Fig. 5B) and spatial registration was performed using the sagittal Allen Brain Reference Atlas (Allen Institute for Brain Science^®^). Confocal image analysis revealed substantial amyloid plaque deposition in the cortex and hippocampus (Fig. 5A) as previously described^27, 31^. Unexpectedly, striatal Aβ plaque density (Fig. 5C) and amyloid plaque number (Fig. 5D, E) were just as high as that seen in the isocortex and hippocampus respectively. Further segmentation of the major brain divisions revealed that amyloid plaque frequency was by far the highest in two subregions, the corpus callosum (11.72% [15/128]) and the caudoputamen (10.93% [14/128], Fig. 5E). Amyloid burden was present in all three functional divisions of the striatum including dorsolateral striatum, dorsomedial striatum and ventral striatum often defined as sensorimotor, associative and limbic striatum respectively^59^. To our knowledge, these results are the first to document the deposition of Aβ in the striatum of middle-aged APP animals, a structure well-established for its role in value-based decision-making and in the learning of reward associations^59, 60^.

**Figure 5.**
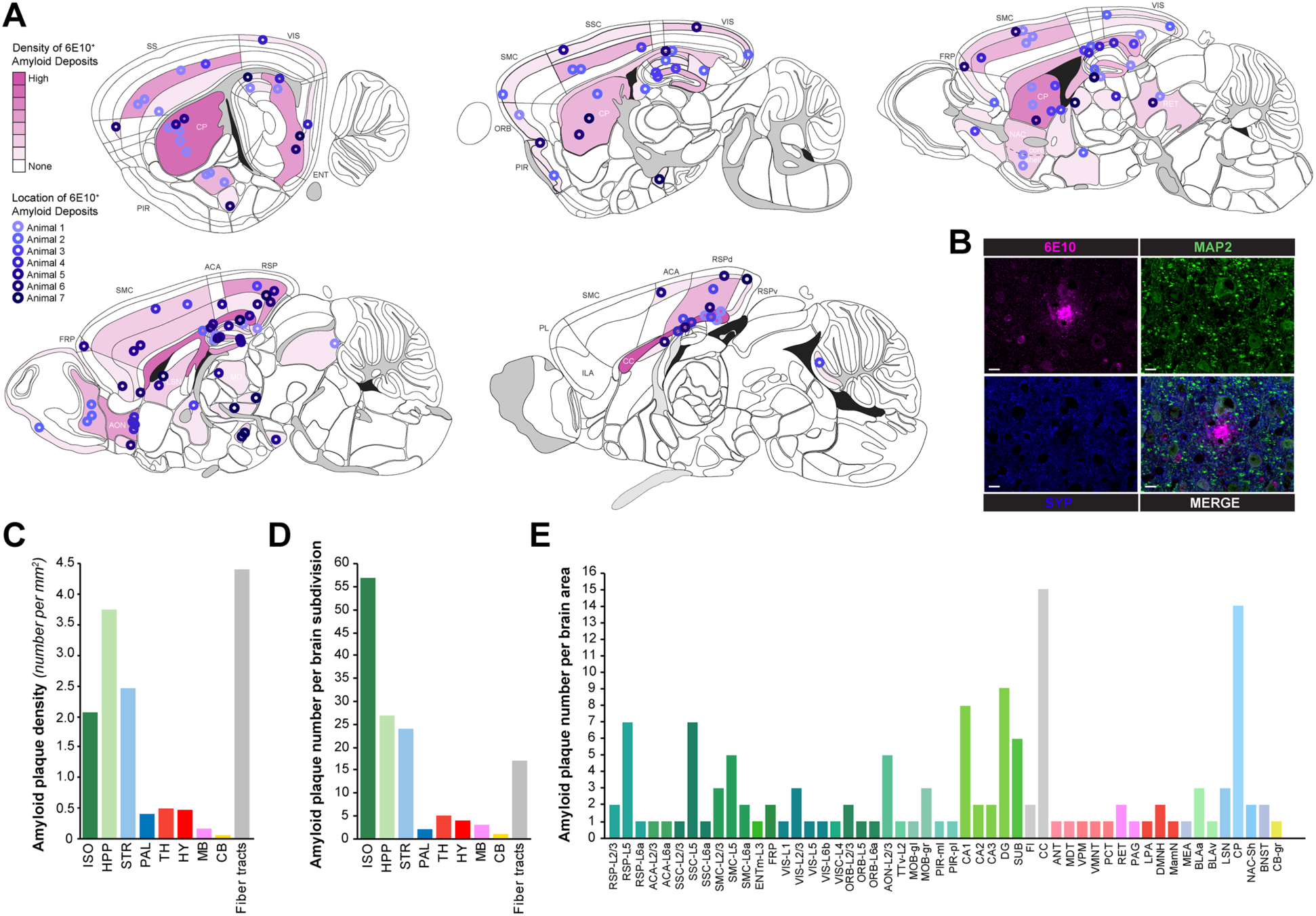
Structure specific localization of amyloid plaques in APP animals. (**A**) Spatial distribution of amyloid plaques throughout the brains of APP animals. Amyloid-β (Aβ) deposits were labeled with 6E10 antibody and confirmed at 40x magnification (blue hues correspond to different animals, pink hues depict relative structure density of plaque load). (**B**) Representative micrograph of an amyloid-β plaque stained for human APP/Aβ (6E10, pink), microtubule-associated protein 2 (MAP2, green), and synaptophysin (SYP, blue) (scale bar = 10 μm). (**C-E**) Quantification of (*C*) amyloid plaque count density in millimeters squared of isocortex (dark green), hippocampus (light green), striatum (light blue), pallidum (blue), thalamus (red-orange), hypothalamus (red), midbrain (light pink), cerebellum (yellow), and fiber tracts (grey), (*D*) total plaque counts per brain region, and (*E*) a breakdown of plaque number by individual structures, referenced to the Allen Brain Atlas.

### Behavioral Differences in APP Mice were Sex Differentiated

The effect of sex is especially important for AD (recently reviewed here^61^) for which phenotype heterogeneity is an intrinsic characteristic. Specifically, women typically present with faster rates of cognitive decline during mild cognitive impairment (MCI) or prodromal AD, and brain atrophy rates are also 1-1.5% faster in women with MCI and AD compared to men^62, 63^. These sex differences are noteworthy considering that putative sex effects on Aβ burden remain unclear in AD. In mice, despite well-established sex differences whereby earlier-onset amyloid pathology has been consistently noted in females across multiple APP transgenic models^26^, sex effects on synaptic or cognitive deficits have not been studied systematically. For these reasons and because previous work from our group only used male mice, we were intrigued to determine whether decision-making processes altered in APP mice were driven by sex.

Following the recommendations made by Shansky^64^, analyses originally compared putative effects of transgene expression between nTG and APP groups but post-hoc data examination also included potential effects of sex differences in both groups of animals. We found significant behavioral differences between males and females in both nTG and APP mice. Male and female nTG mice looked similar in many behaviors, with the exceptions of laps run and WZs quit (Suppl. Figure 7). Female nTG mice generally ran more laps (RM-ANOVA stage 3: *F*_(1,13)_ = 9.287, *p* = 0.005; epoch A: *F*_(1,13)_ = 20.27, *p* < 0.0001; epoch B: *F*_(1,13)_ = 22.46, *p* < 0.0001; epoch C: *F*_(1,13)_ = 12.38, *p* = 0.0006; Suppl. Figure 7A). Female nTG mice also quit more WZs after entering (RM-ANOVA stage 3: *F*_(1,13)_ = 4.23, *p* = 0.05; epoch A: *F*_(1,13)_ = 21.42, p < 0.0001; epoch B: *F*_(1,13)_ = 10.55, *p* = 0.001; Suppl. Figure 7E).

We also found significant sex differences amongst APP mice (Suppl. Figure 8). Generally, male and female APP mice ran an equivalent number of laps and earned the same amount of pellets, with males earning slightly more in the middle of the last epoch (RM-ANOVA epoch B: F_(1,13)_ = 8.17, *p* = 0.005; Suppl. Figure 8B) and females earning more pellets at the end of the experiment (RM-ANOVA days 53-70: *F*_(1,13)_ = 8.86, *p* = 0.003; Suppl. Figure 8B). This corresponded with males having a shorter IEI during the middle of the last stage (RM-ANOVA epoch B: *F*_(1,13)_ = 4.27, *p* = 0.04; Suppl. Figure 8C) and females having a shorter IEI at the end (RM-ANOVA epoch C: *F*_(1,13)_ = 12.09, *p* = 0.0008; Suppl. Figure 8C).

More striking behavioral sex differences were found in the decisions to enter or skip offers. Female APP mice entered, and therefore accepted more offers (RM-ANOVA epoch B: *F*_(1,13)_ = 57.92, *p* < 0.0001; epoch C: *F*_(1,13)_ = 7.86, *p* = 0.006; Suppl. Figure 8D) as well as quit more WZs (RM-ANOVA stage 2: *F*_(1,13)_ = 8.68, *p* = 0.007; stage 3: *F*_(1,13)_ = 11.59, *p* = 0.002; epoch A: *F*_(1,13)_ = 18.2, *p* = 0.0003; epoch B: *F*_(1,13)_ = 175.94, *p* < 0.0001; epoch C: *F*_(1,13)_ = 82.53, *p* < 0.0001; Suppl. Figure 8D). Due to female APP mice entering more offers than their male based counterparts, female APP mice had higher OZ thresholds (RM-ANOVA epoch B: *F*_(1,13)_ = 41.93, *p* < 0.0001; epoch C: *F*_(1,13)_ = 10.33, *p* = 0.002; Suppl. Figure 8E). Together this data suggest that male APP mice were more selective about the offers they entered (by entering significantly fewer WZ than females) and less likely to quit upon entering. By contrast, the OZ and WZ threshold patterns displayed by female APP mice were reminiscent of those generally observed in nTG controls.

Given that female APP mice looked more similar to nTG mice than male APP mice, we examined whether sex affected choice deliberation differentially in APP animals. When examining VTE behavior in the OZ, male APP mice had higher VTE values than female APP mice (RM-ANOVA epoch B (days 23-52): *F*_(1,29)_ = 80.05, *p* < 0.0001; epoch C (days 53-70): *F*_(1,29)_ = 49.61, *p* < 0.0001; Fig. 6A) when deciding to eventually enter. When VTE for both entering and skipping were averaged over the last 10 days of the experiment, when animals are well trained, female APP mice showed sharp distinctions in the amount of VTE demonstrated when entering versus skipping, looking similar to nTG mice, with higher VTE for skipping than entering (one-way ANOVA, *F*_(1,4)_ = 232.7, *p* = 0.0001). Males on the other hand showed similar levels of VTE whether entering or skipping (*F*_(1,6)_ = 0.49, *p* = 0.51). Looking at VTE based on value, female APP mice looked more similar to control mice again, deliberating more on bad offers than good (one-way ANOVA, *F*_(1,4)_ = 33.27, *p* = 0.004; Fig. 6C, D). Again, male APP mice showed the same deliberation for good and bad deals (one-way ANOVA, *F*_(1,6)_ = 0.39, *p* = 0.56; Fig. 6C, D). Finally, we examined the VTE levels by flavor preference to see if there were sex differences here as well. As a reminder, nTG mice showed higher VTE for their least preferred flavors, showing lower VTE (and smooth entrances) for their most preferred. APP mice showed high VTE values, regardless of flavor preference. Here again, females looked more similar to nTG mice, showing distinct differences in VTE depending on flavor beginning in the second stage of the experiment (RM-ANOVA first: *F*_(3,56)_ =1.31, *p* = 0.28; second: *F*_(3,40)_ = 7.32, *p* = 0.0005; third: *F*_(3,40)_ = 5.36, *p* = 0.003; epoch A: *F*_(3,40)_ = 5.57, *p* = 0.003; epoch B: *F*_(3,240)_ = 11.12, *p* < 0.0001; epoch C: *F*_(3,144)_ = 4.47, *p* = 0.005; Suppl. Figure 6F). Unlike their female counterparts, male APP mice showed distinct differences in the amount of VTE they used for flavor preferences, at any point in the experiment (RM-ANOVA, first: *F*_(3,80)_ = 0.38, *p* = 0.77; second: *F*_(3,60)_ = 1.44, *p* = 0.24; third: *F*_(3,60)_ = 2.1, *p* = 0.11; epoch A: *F*_(3,60)_ = 0.78; *p* = 0.51; epoch B: *F*_(3,360)_ = 1.01, *p* = 0.39; epoch C: *F*_(3,216)_ = 1.11, *p* = 0.31; Suppl. Figure 6D).

**Figure 6.**
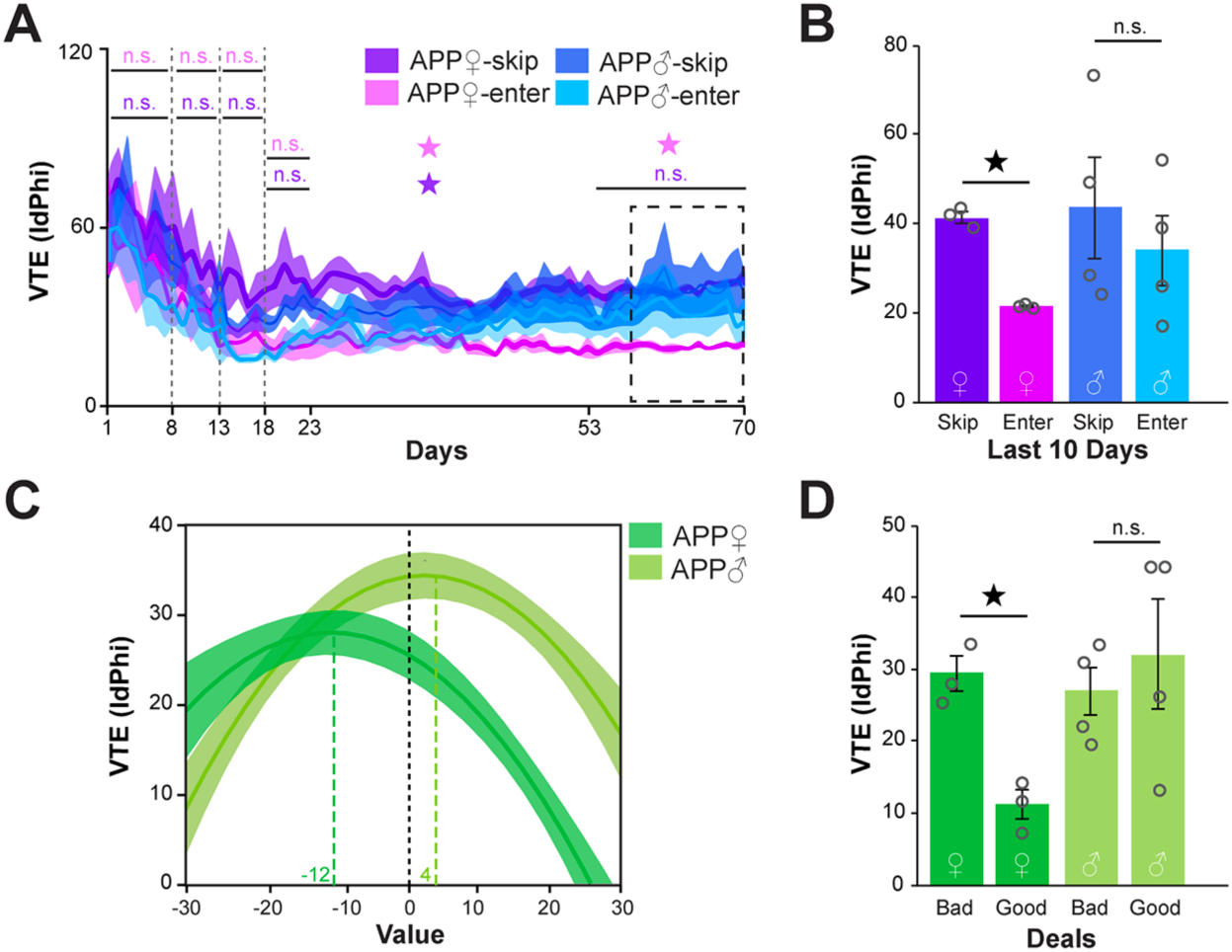
Vicarious-trial-and-error split by sex. (**A**) Average VTE (*IdPhi*) values split by skip versus enter decisions across days of training in APP mice split by sex (blue = male APP, purple and pine = female APP). Open star denotes p<0.05 for between-group differences in skip VTE. Closed star denotes p <0.05 for between group differences in enter VTE. Data are presented as the daily means (± SEM) across the entire experiment. Vertical dashed lines represent stage transitions. (**B**) Bar graph depicting mean ± SEM for well-trained animals (last 10 days) to illustrate sex differences. *, p<0.05 one-way ANOVA within group (female APP). (**C**) Quadratic functions of VTE by value (WZ Th. minus offer) for the last 10 days. Negative value denotes an economically unfavorable offer. Peaks are significantly right-shifted for male APP mice (value of +4 compared to −12 for female APP mice), indicating higher VTE for closer to neutral values. (**D**) Mean ± SEM of values from E for bad (value <0) and good (value >0) deals *, p<0.05 one-way ANOVA within group (female APP mice).

### Spatial differences in Aβ pathology are sex specific in APP mice

Upon identifying sex-dependent alterations in decision-making from APP mice performing the RRow task, we wondered whether amyloid burden was also influenced by sex differences. Unlike previous studies which reported enhanced amyloid deposition in female animals from several other APP transgenic mouse models^26^, averaged plaque numbers did not differ between APP male and female mice (*t* test, *p* = 0.424, Suppl. Figure 9A). However, factoring in the localization of Aβ deposits revealed that the proportion of amyloid plaques per brain were different between the sexes (Suppl. Figure 9B-D). Notably, inverse proportions of plaque deposition were observed in male APP mice compared to female APP littermates in isocortical and hippocampal divisions whereby Aβ deposits were proportionally more abundant in the isocortex of female animals (33 in female vs. 27 in male mice respectively) but less abundant by a factor of two-fold in their hippocampi (9 in female vs. 18 in male mice respectively; Suppl. Figure 9C). This observation was further supported upon increasing segmentation of the brain subdivisions by areas and layers, when applicable (Suppl. Figure 9D). Out of 27 cortical areas/layers, amyloid plaques were detected in 20 specific locales from female APP brains (i.e., 74.1%) whereas only 11 locales displayed Aβ deposits in male APP mice (i.e., 40.7%). In sharp contrast, amyloid plaques were not only found in all hippocampal fields from male APP brains unlike in female APP mice, their relative proportions within these locales were also quantitatively much higher as exemplified by the 80/20 and 66/33 ratios observed for CA1 and subiculum respectively (Suppl. Figure 9D). This novel sex-specific finding is particularly interesting considering that both of these hippocampal domains critically determine diverse behavioral and cognitive functions.

### Striatal inhibitory network alterations in male APP mice

Because network activities supporting cognition are altered prior to disease onset in AD and because interneuron dysfunction has emerged as a potential mechanism underlying these network abnormalities (see for review^65^), we measured the ectopic expression of neuropeptide Y (NPY), a well-established marker of molecular alterations linked to the network remodeling in APP mice^27, 32, 66–68^, in all mice subjected to RRow. Adapting the approach developed by the Mucke group^66^ for confocal imaging analysis, we observed a ∼2-fold increase in NPY immunoreactivity in mossy fiber axons in the stratum lucidum (SL) of APP mice (*t* test, *F*_(1,15)_ = 3.5321, *p* = 0.0414; Suppl. Fig. 10A,B). A more modest elevation in NPY immunoreactivity was also found in the stratum lacunosum moleculare (SLM) of APP mice (*t* test, *F*_(1,15)_ = 4.4108, *p* = 0.0279; Suppl. Fig. 10A,C). In both subfields, putative sex effects were not found for SL axons (two-way ANOVA, *F*_(3,15)_ = 1.8615, *p* = 0.1945; Suppl. Figure 10D) or SLM axons (two-way ANOVA, *F*_(3,15)_ = 2.6094, *p* = 0.1041; Suppl. Figure 10E). These hippocampal findings are consistent with prior observations from 7- to 10-month-old APP animals^32^.

Since Aβ pathology was newly identified in the striatum of APP mice, we also evaluated the ectopic expression of NPY in the caudoputamen and nucleus accumbens (Fig. 7A). When comparing nTG and APP mice, NPY immunoreactivity was similar across genotype groups (*t* test, *F*_(1,15)_ = 1.8185, *p* = 0.2009 and *F*_(1,15)_ = 2.1196, *p* = 0.1711 respectively; Fig. 7B,C). However, considering sex as a variable revealed sex-specific alterations in NPY expression whereby male nTG mice displayed high levels of ectopic NPY expression in the caudoputamen which was not present in male APP animals (two-way ANOVA followed by Tukey HSD, *F*_(3,15)_ = 9.3868, *P* = 0.0023 with an effect of transgene *F*_(1,15)_ = 5.0201, *P* = 0.0406, sex *F*_(1,15)_ = 13.5644, *P* = 0.0036, and transgene*sex interaction *F*_(1,15)_ = 8.0451, *P* = 0.0162). In addition, these basal high amounts of caudoputamen NPY expression in male nTG mice was not observed in female nTG mice (Fig. 7D), consistent with a previous report detailing a sex-specific difference of striatal NPY expression in rats (70). Despite falling just short of statistical significance, similar trends were observed in the nucleus accumbens (two-way ANOVA followed by Tukey HSD, *F*_(3,14)_ = 3.6874, *p* = 0.0507). Altogether, these results suggest that Aβ pathology is associated with alterations in network activity in the striatum of APP mice.

**Figure 7.**
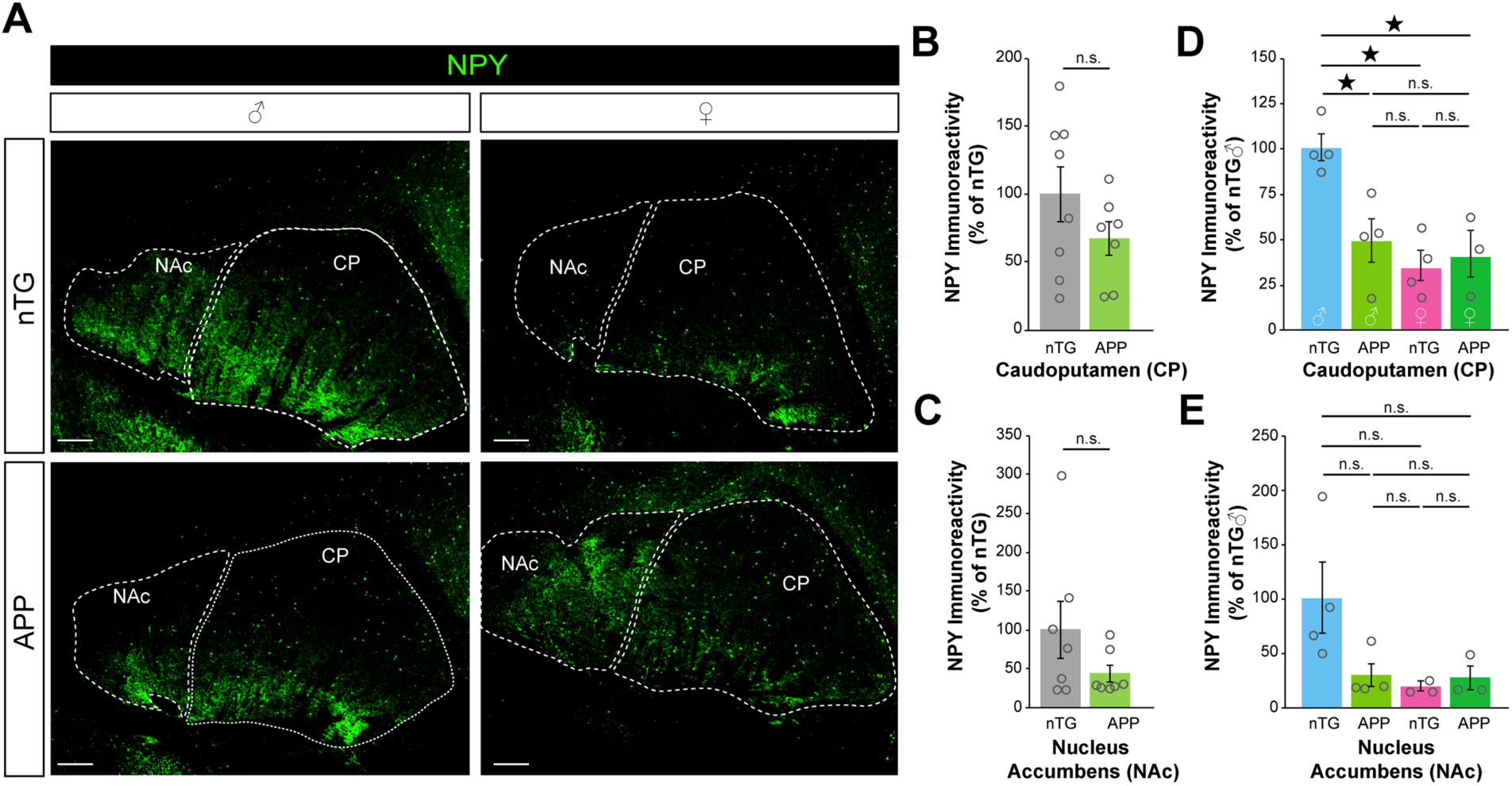
Sex-specific loss of striatal NPY in J20 mice. (**A**) Representative immunostaining of the striatum from nTG and APP male and female animals stained for neuropeptide-y (NPY, green). (**B,C**) Quantification of NPY immunoreactivity in the (*B*) caudoputamen (CP) and (*C*) nucleus accumbens (NAc) of nTG and APP mice (nTg n = 8, APP n = 7, CP *t* test, F_(1,15)_ = 1.8185, P = 0.2009, SLM *t* test, F_(1,15)_ = 2.1196, P = 0.1711). (**D,E**) Quantification of NPY immunoreactivity in the (*D*) CP and (*E*) NAc in the same cohort with groups divided by sex (nTG male n = 4, APP male n = 4, nTG female n = 4, APP female n = 3, CP two-way ANOVA followed by Tukey HSD, F_(3,15)_ = 9.3868, P = 0.0023 with an effect of transgene F_(1,15)_ = 5.0201, P = 0.0406, sex F_(1,15)_ = 13.5644, P = 0.0036, and transgene*sex interaction F_(1,15)_ = 8.0451, P = 0.0162; NAc two-way ANOVA followed by Tukey HSD, F_(3,14)_ = 3.6874, P = 0.0507). Data are presented as mean ± SEM.

## DISCUSSION

To better understand how complex decision-making may be altered with early stages of Alzheimer’s disease, we subjected a widely used mouse model of AD, the J20 line, and young adult nTG littermates to a neuroeconomic spatial foraging task called RRow that accesses multiple decision-making systems. We found that the strategies used by APP mice on this task differed drastically from that of the non-transgenic controls^43^, with APP animals relying heavily on VTE events and showing impaired procedural decision-making. These behavioral differences were largely accounted for by the male APP mice, with female APP mice behaving similarly to nTG mice. Neuropathological analyses of Aβ deposits throughout the whole brain unexpectedly revealed that 9-month-old APP mice displayed much more widespread plaque aggregation than has been reported previously. This was particularly notable in the striatum, which has never been reported. Along with this plaque deposition, ectopic NPY expression was significantly decreased in the striatum of male APP mice suggesting network remodeling of this region.

Theories of decision-making suggest that there are at least three dissociable systems: a Pavlovian system that chooses unconditioned responses based on associations between stimuli and outcomes^16, 40^, a procedural system that chooses actions based on learned associations between actions and stimuli^16, 39^, and a deliberative system that considers how an action influences future possibilities^15, 17, 35^. Vicarious-trial-and-error behavior has been shown to be indicative of deliberation^58^. During VTE, in rats, hippocampal representations sweep forward, alternating potential goals that are synchronized with reward value representations suggesting that outcome predictions are being evaluated^58, 69, 70^. Control mice in this task showed higher VTE when eventually deciding to skip an offer versus deciding to accept an offer, suggesting that the decision to accept an offer, particularly a good offer, likely arises from a non-deliberative decision-system (either Pavlovian or procedural). This discrimination replicates previous observations of mice on this task^43^. Additionally, nTG mice showed higher VTE when presented with bad offers (value below their threshold to wait; see methods) and with their least preferred flavor of food. In those cases, deliberative decision-making is used to counter instinctual approach behaviors. In stark contrast to the distinction nTG mice demonstrated, APP mice showed equal amounts of VTE whether deciding to skip or enter, suggesting deliberative mechanisms of decision-making were paramount in both taking and rejecting offers. Additionally, APP mice deliberated equally for both good and bad offers (high or low values), as well as for their most and least preferred flavors, which was fundamentally different than the nTG control mice.

Our observations largely suggest that APP mice rely excessively on deliberation even in situations where nTG mice use procedural systems. One hypothesis for the excessive deliberation observed in the APP mice may reflect an inability to process the planning signals normally generated by the hippocampus, potentially due to the amyloid pathology in the hippocampus. Another potential hypothesis for this high deliberative behavior may be that the amyloid pathology in the striatum disrupted procedural decision making, forcing reliance on the deliberative system. Evidence largely supports the later hypothesis, as APP mice actually normalize their behavior faster than nTG controls. This suggests that the excessive deliberation observed was functional and helpful to their processing. This disrupted procedural decision-making hypothesis is further supported by the sex-linked differences and changes in NPY functionality observed in the striatum.

As RRow gives us the ability to examine multiple forms of decision making that encompass multiple brain regions, it was of critical importance to us to possess a broad characterization of amyloid pathology throughout the brain. Previous work has extensively examined amyloid plaques in the hippocampus^27^ throughout the lifespan of APP mice but rare studies have determined amyloid burden across the brain^33^. The mice used in our study were 9-months of age at the time of tissue processing and showed significant pathology in the hippocampus, as was expected. What was not expected was how dense plaque aggregation was in the striatum. In these animals, the striatum showed the second highest plaque density. The fact that both the hippocampus and the striatum had high plaque deposition suggests that neuronal functioning may have been impaired in striatum as well as, or even more than, the hippocampus. It is well established that the hippocampus is centrally implicated in spatial navigation and memory^15, 35^. And though proper lateral striatal functioning is known for being important for procedural decision making^71^, the hippocampus and the medial striatum likely work in concert to switch between rigid and flexible behaviors through connections between the prefrontal cortex^13, 34, 72^. Thus, the fact that both the hippocampus and the striatum showed plaque deposits suggests that the impairment in procedural decision-making displayed by APP mice could result from effects of pathology, perhaps on either procedural decision systems or on decision-system-conflict mediation systems.

It has recently emerged that including imaging data from subcortical areas might be critical for the clinical presentation of AD. Supporting this view, several studies reported that high striatal Aβ burden predicts faster cognitive decline than high cortical Aβ loads in humans with mild cognitive impairment (MCI) or AD^22, 23^. It is important to note that despite this advancement, the impact of these striatal amyloid plaques on human cognition are currently unknown and requires further studies. The neuropathological findings of our study documents large depositions of Aβ plaques in the striatum of APP mice, reminiscent of the striatal Aβ plaques recently reported in subjects with AD^22, 23^. This parallel is even more striking considering that familial AD is distinguished from late-onset AD by early striatal Aβ deposition and considering that APP mice like J20 animals harbor FAD mutations^23, 27^. The novel description that Aβ deposition in the striatum of APP mice is associated with abnormal alterations of procedural decision-making provides support for an impact of striatal Aβ pathology on behavior for the first time in this model. Because RRow has been successfully translated to a similar neuroeconomic task called WebSurf for use in human subjects^44^, our studies using preclinical models of AD open up the possibility of testing patients with prodromal AD or MCI using this approach.

Because we included both males and females in our experimental design, we conducted post-hoc analyses to investigate any potential sex differences in the behavior we observed. Epidemiologic studies of AD are reported that women typically present with a more rapid cognitive decline^62^ while imaging studies indicated that women also present with an exacerbated brain atrophy when compared to men^63^. These observations and others indicate that sex is an important contributor of disease heterogeneity^61^, although the causes underlying these potential differences are unclear. In most mouse models of AD, female mice have been shown to have Aβ pathology develop earlier than male mice^26^. In our studies, the most striking behavioral differences between nTG and APP mice, namely high reliance on deliberative behavior and impaired procedural decision-making in APP mice, were largely carried by male APP animals. In fact, in most cases, female APP mice looked very similar to control nTG littermate animals. Although the average plaque number was similar between males and females, segregating by subregions revealed that males carried a higher plaque burden in the hippocampus, whereas females carried the largest Aβ burden in the isocortical regions. It is thus possible that the sex differences in observed behaviors are due to this differential localization of Aβ deposits. Further studies will be needed to rigorously establish causal links.

These data also highlight the need for sensitive behavioral paradigms that allow for the dissociation of multiple valuation processes which is difficult using standard tasks that have been previously used to characterize APP mice. For instance, APP mice of the same age as used in this study are characterized as having impaired spatial reference memory. This has been assessed using standard spatial memory tasks, such as the Barnes maze and the Morris water mazes^54, 57^. Duration to find the platform or hole and the distance used to find the platform/hole are the primary quantitative measures for cognitive memory performance in these tasks. A more detailed analysis of behaviors used when searching for the goal allows for a finer dissection of neural circuits involved and elucidates different strategies that may be used that are obscured by measurements of only distance and speed. Recent examples of groups using the Morris water maze have discovered different search strategies between transgenic and non-transgenic mice^73^. These fine analyses allow for group differences to be revealed even if both groups look equivalent in other more commonly reported measures such as distance traveled or latency to find the platform. Without careful examination and dissection of the finer details of behavior, important findings are obscured. When only considering earns in our task, it would have been easy to just say that APP mice did better at this task than nTG mice or that they learned the task faster. What is lost in limited analyses would be the very disparate strategies used by both groups in response to a changing economic environment.

In conclusion, APP mice displayed impaired procedural decision-making and that, to compensate for this impairment, these animals relied on deliberation to succeed in a neuroeconomic decision-making task. To our knowledge, this is the first study to report a previously undocumented deposition of Aβ in the striatum of J20 mouse line, which is associated with aberrant ectopic expression of NPY and sex-specific alterations in decision-making while performing a neuroeconomic task. These studies are directly relevant to the recently described striatal deposition of Aβ in FAD carriers and may extend to subjects with late-onset AD.

## MATERIALS AND METHODS

### Mice

Eight transgene-positive (4 male and 4 female) and eight transgene-negative J20-C57BL/6J APP transgenic mice (4 male and 4 female; The Jackson Laboratory, #006293) all 6-months of age were used for this study. During the experimental procedure, one female J20 mouse became sick and was eliminated from the study, leaving N_J20_=7. J20 mice express a mutant form of human APP by way of the Swedish (KM670/671NL) and Indiana (V717F) mutations, resulting in higher levels of total human amyloid-beta and an increase in the Aβ42/Aβ40 ratio, respectively. This transgene is driven by the PDGF-β promoter. Mice were single-housed (beginning at 6 months) in a temperature- and humidity-controlled environment with a 12 h light/12 h dark cycle with water ad libitum. Mice were food restricted to 90% free-feeding body weight and trained to earn their entire day’s food ration during their 1-hour Restaurant Row session. All experiments were approved by the University of Minnesota Institutional Animal Care and Use Committee (IACUC) and adhered to NIH guidelines. All mice were tested at the same time (beginning at 8:30 am) every day during their light phase in a dimly lit room. Mice were weighed before and after each session to make sure they were above 90% of their free-feeding body weight and were fed a small post-session ration of food (1.5-2g) on the occasion that their body weight fell below the 90% guideline.

### Pellet training

One week prior to the start of training on Restaurant Row, mice were trained to eat the pellets that were used in the task. During this time, mice were taken off their regular food and introduced to a single daily serving of BioServ full-nutrition 20 mg dustless precision pellets (5 g). This serving consisted of an equal mixture of all four flavors found in the maze; chocolate, banana, grape, plain. One day prior to the start of training, mice deprived of their previous day’s ration were introduced to the maze. Each mouse was given 20 minutes to explore the maze and familiarize themselves with the feeding sites which were filled with excess food of the particular flavor that would be found during the experiment. Each restaurant location was marked with unique spatial cues and remained constant throughout the entire experiment.

### Restaurant Row

In this task, food deprived mice are allowed one hour to traverse a square maze with four feeding sites that offer different flavors of food at varying delays (i.e., costs). Mice need to learn to balance their food preferences against the potential cost at each reward-site encounter in order to obtain their only source of food for the day. Because mice have limited time to forage on the task, the delays that they must wait for food are analogous to costs spent from a limited (time) budget. As mice progress through the stages of learning, the range of delays increases and thus the reward environment grows increasingly scarce. As rewards become scarce, conflicts in decision-making arise forcing the animals to adapt new foraging strategies that may no longer suffice in previously rich environments.

Each daily session lasted one hour. At the beginning of the test, one restaurant was randomly selected to be the starting restaurant. An offer was made if mice entered the restaurant’s offer zone **(OZ**) from the appropriate direction in a counterclockwise manner. An offer began when the mouse entered the OZ and consisted of a delay that the mouse would need to wait before earning a pellet upon entering the wait zone (**WZ**). Brief tones (4,000-15,223 Hz for 500ms followed by 500ms of silence) sounded upon entry into the OZ, with pitch indicating the delay of the offer.

Tones repeated every second until mice either *left* the OZ for the next restaurant (**skip**) or *entered* the WZ (**enter**). Upon entering the WZ, the tones counted down (in 387 Hz steps) each second until the mouse either *left* (and **quit**) the WZ, or the countdown reached 0 (following the final 1s tone = 4,000 Hz), at which point a pellet was dispensed (**earn**). If the mouse left (quit) the WZ during the countdown, the tone stopped, and the offer was rescinded. To discourage mice from hoarding earned pellets, motorized feeding bowls cleared uneaten pellets after the mouse exited the WZ. Mice quickly learned not to leave the WZ without consuming the earned pellet. The next restaurant in the counterclockwise sequence was always and only the next available restaurant where an offer could be made. This ensured mice encountered offers across all restaurants in a fixed consecutive order.

Training was broken into four stages. During the first stage (days 1-7), mice were given only 1 sec offers for all restaurants. During the second stage (days 8-12), mice were given offers that ranged from 1 to 5 sec (4,000 Hz to 5,548 Hz in 387 Hz steps). Offers were pseudo-randomly selected, such that all 5 offer lengths were encountered in 5 serial trials before being reshuffled, ensuring a uniform distribution of offer lengths. Stage 3 (days 13-17) consisted of offers from 1-15 s (4,000-9,418 Hz). Stage 4, the final stage (days 18-70), consisted of offers ranging from 1-30 s (4,000-15,223). Again, all offers in all stages were pseudo-randomly selected in each restaurant independently and all offers were encountered before being reshuffled.

To assess flavor preferences, the total earnings of each flavor at the end of the session were examined. Flavors were ranked from most earned to least earned for each individual mouse. Flavor preferences were established by the beginning of the second stage and remained consistent throughout the rest of the experiment.

Four Audiotek tweeters positioned next to each restaurant were powered by Lepy amplifiers to play tones at 70 dB in each restaurant. Med Associates 20 mg feeder pellet dispensers and 3D-printed feeder bowls fashioned with mini-servos to control automated clearance of uneaten pellets were used for pellet delivery. Animal tracking, task programming, and maze operation were powered by AnyMaze (Stoelting).

### Immunohistochemistry

The left hemisphere of each animal’s brain was sent to NeuroScience Associates for MultiBrain® processing. Upon return, matrix sheets of 30 μm-thick sections were triple-labeled for Synaptophysin (1:1500, Synaptic Systems 101004), 6E10 (1:200, BioLegend 803002), and MAP2 (1:400, Novus NB300-213). Secondaries used were Alexa Fluor 488, 555, and 647 (1:400, Invitrogen A-11073, A-21127, A32933). Sections were treated with 0.1% Sudan Black for auto-fluorescence and mounted with Prolong Gold mounting media (Invitrogen P36930). Confocal imaging was done using Olympus FluoView FV1000. Immunostained hemibrain sections were acquired by 4X tiling and each amyloid plaque was further confirmed at 40X magnification. For plaque spatial registration, the sagittal Allen Adult Mouse Brain Atlas (<u>atlas.brain-map.org</u>) was used as a reference.

### Quantification of Inhibitory Network Activity

To investigate inhibitory network activity, we followed previously described protocols for quantification of NPY immunofluorescence^66^. In brief, sagittal sections embedded and sectioned by NeuroScience Associates were stained with anti-neuropeptide-y (Cell Signaling 11976, 1:400), goat anti-rabbit Alexa Fluor 555 Plus (ThermoFisher A32732, 1:400), MAP2, goat anti-chicken Alexa Fluor 647 Plus, and imaged using an Olympus FV1000 microscope. Hippocampal sections were sequentially acquired using a 10x objective with NA of 0.4 and striatal sections were sequentially acquired using a 4x objective with NA of 0.16. For imaging both structures, the Alexa Fluor 555 Plus was excited at 559nm and 10.0% transmissivity and emissions were collected from 575-620nm with a Kalman integration of 10. Maximum grey values were lowered in FIJI to match the upper end of the distribution peak. The minimum grey value was increased until the background level measured in the granule layer of the dentate gyrus for the hippocampus or the corpus callosum for the striatum was consistent across all images of the same structure. For quantification of NPY expression in the hippocampus, regions of interest were selected over the stratum lacunosum moleculare (SLM) and the stratum lucidum (SL). In the striatum, the selected regions of interest were the caudoputamen (CP) and the nucleus accumbens (NAc). All regions of interests were drawn over the MAP2 channel with the help of the Allen Brain Atlas. The calculated mean grey values of the regions of interest were normalized to their structure’s respective background control area that was used to determine the minimum grey value. In addition, two sections for each animal were stained and used to calculate average values for the regions of interest.

### Statistical analysis

All data were processed in Matlab and statistical analyses were carried out using JMP Pro 13 Statistical Discovery software package from SAS. All data are expressed as mean +/− SEM. Offer zone thresholds were calculated by fitting a sigmoid function to offer zone choice outcome (skip versus enter) as a function of offer length for all trials in a single restaurant for a single session and measuring the inflection point. Wait zone thresholds were calculated by fitting a sigmoid function to wait zone outcomes (quit versus earn) as a function of offer length for all entered trials in a single restaurant for a single session. For analyses that depend on thresholds, analyses at each timepoint used that specific timepoint’s threshold information. Statistical significance was assessed using Student *t* tests, one-way, two-way, and repeated measures (RM) ANOVAs, using mouse as a random effect in a mixed model, with post-hoc Tukey *t* tests correcting for multiple comparisons. Significance testing of immediate changes at block transitions within group were tested using repeated measures ANOVA between 1 d pre- and 1 d post-transition. These are indicated by significance annotations on the dotted lines denoting transitions on relevant figures. Significance testing of behavior differences between groups were tested using a repeated measures ANOVA across all days within a given block. These are indicated by significance annotations *within* the plot. The period of renormalization was estimated based on animal driven performance improvements in the 1-30 s stage and not imposed on the animals by experimenters nor the protocol design. Renormalization was characterized by identifying the number of days in the 1-30 s block, after which total pellet earnings and reinforcement rate reliably stabilized and was no different from performance in relatively reward-rich environments in earlier stages of the experiment.

## Abbreviations

AD: Alzheimer’s disease
NTG: non-transgenic
APP: amyloid precursor protein
OZ: offer zone
WZ: wait zone
VTE: vicarious trial and error
RT: reaction time
IEI: inter-earn-interval

## Acknowledgements & Funding Sources

This work was supported by grants from the National Institutes of Health (NIH) to SEL (RF1-AG044342, R21-AG065693, R01-NS092918, R01-AG062135 and R56-NS113549), to ADR (R01-MH080318, R01-MH112688) and to MJT (R01-DA041808). Additional support included start-up funds from the University of Minnesota Foundation and bridge funds from the Institute of Translational Neuroscience to SEL.

## Data availability statement

The datasets generated during and/or analyzed during the current study will be available on Dryad.

## Supplementary Figure legends

**Supplementary Figure 1.**
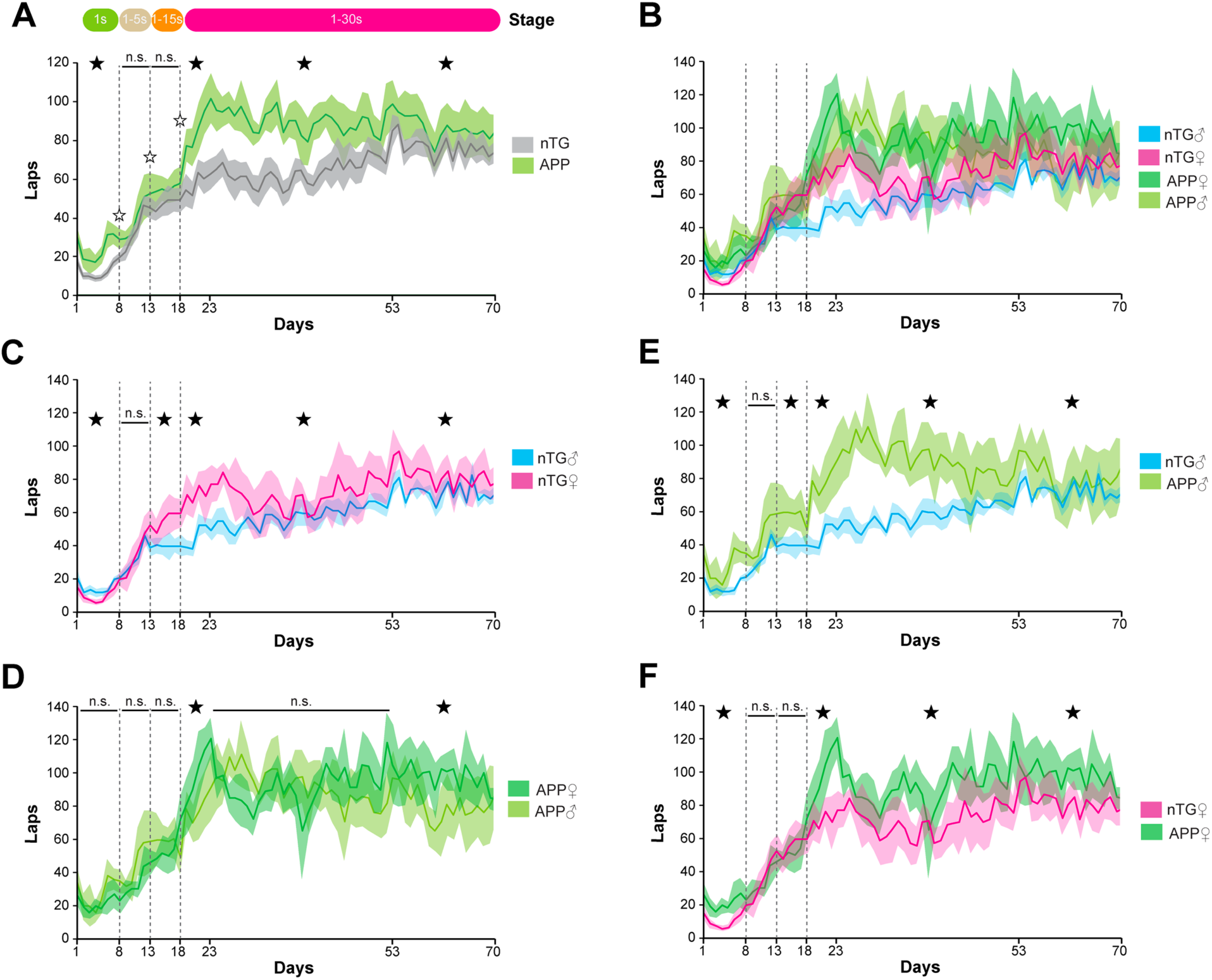
Laps run in the correct counterclockwise direction. (**A**) Laps run between groups. APP mice run more laps than nTG mice. (**B**) Laps in groups split by sex. (**C**) Laps run by nTG mice split by sex. Female nTG mice run more laps than male nTG mice. (**D**) Laps run by APP mice split by sex. (**E**) Laps run by male mice split by group. Male APP mice run significantly more laps than male nTG mice. (**F**) Laps run by female mice split by group. Female APP mice run more laps than female nTG mice. Data are presented as the daily means (± SEM) across the entire experiment. Vertical dashed lines represent stage transitions.

**Supplementary Figure 2.**
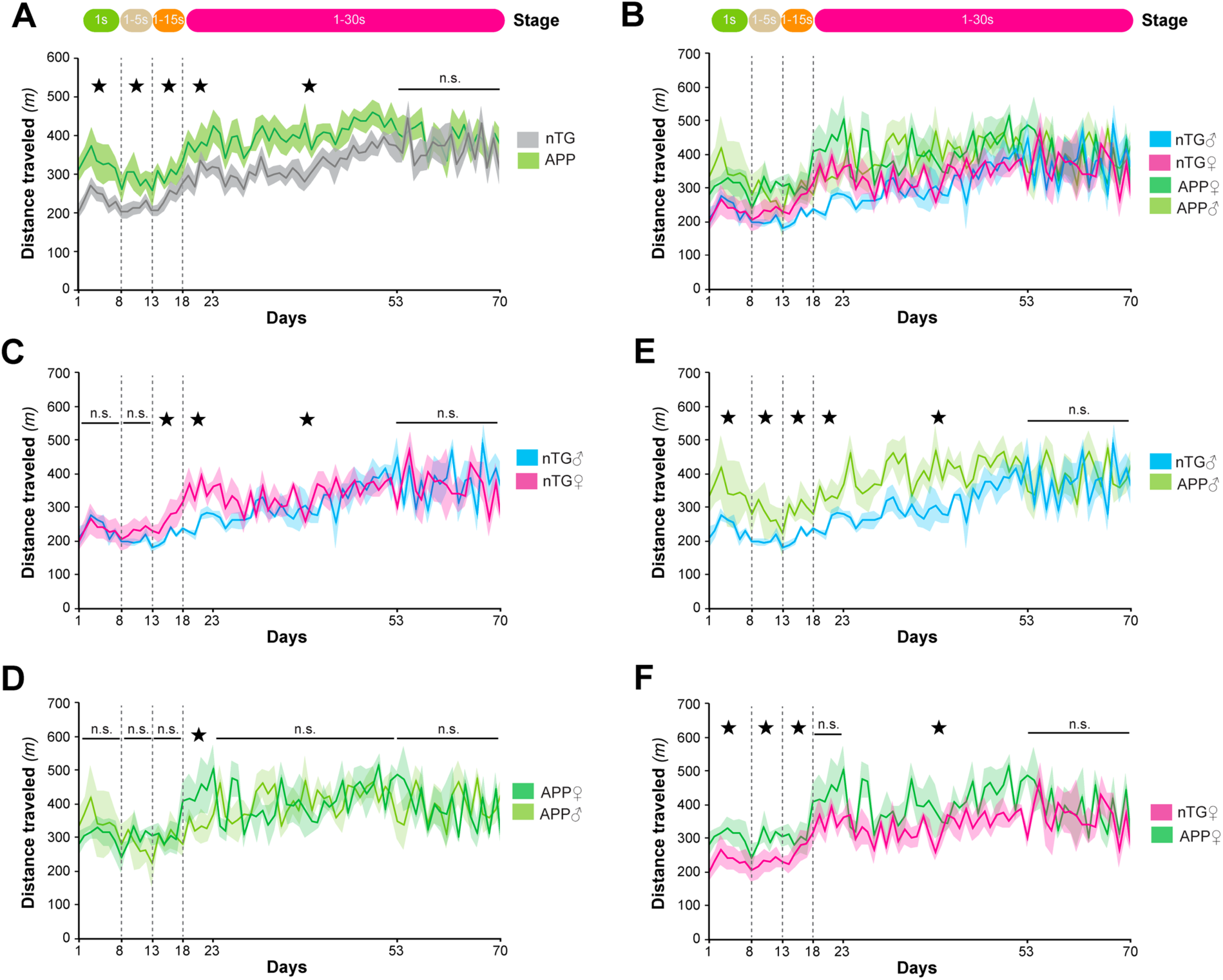
Distance traveled during the 1-hour experimental sessions. (**A**) Distance travelled in meters (m) across groups. APP mice run significantly further during their 1- hour session until the last epoch during stage 3 (days 53-70). (**B**) Distance travelled by groups split by sex. (**C**) Distance traveled by nTG mice split by sex. (**D**) Distance travelled by APP mice split by sex. (**E**) Distance travelled by male mice split by groups. Male APP mice run significantly further than male nTG mice at every stage of the experiment except for the last epoch (days 53-70). (**F**) Distance travelled by female animals split by groups. Female APP mice run significantly further than female nTG mice at every stage of the experiment except for the last epoch (days 53-70). Data are presented as the daily means (± SEM) across the entire experiment. Vertical dashed lines represent stage transitions.

**Supplementary Figure 3.**
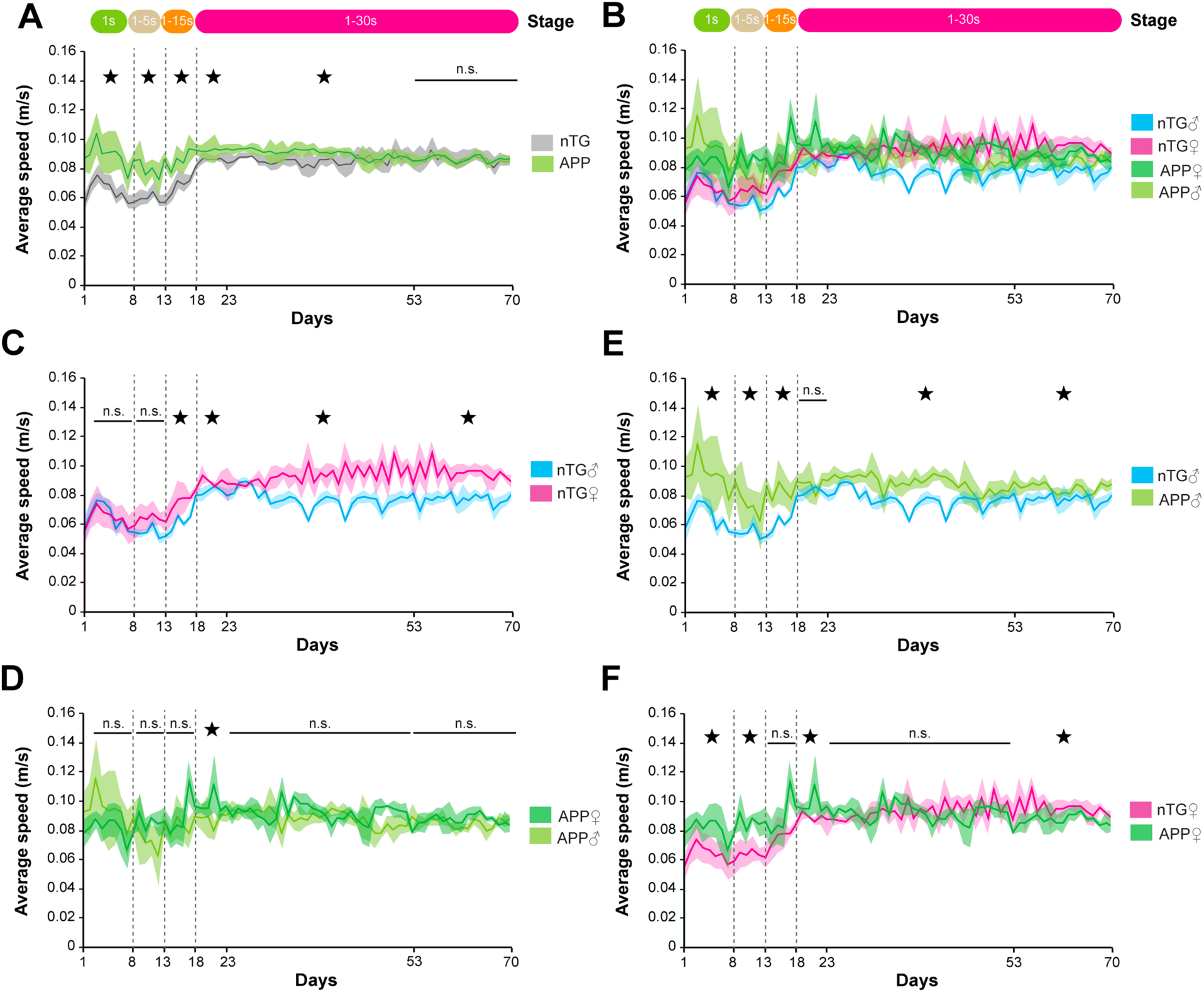
Average speeds during the 1-hour experimental sessions. (**A**) Average speed (m/s) across mouse groups. APP mice run significantly faster during the 1-hour session until the last epoch during stage 3 (days 53-70). (**B**) Average speeds by groups split by sex. (**C**) Average speeds by nTG mice split by sex. Female nTG mice run faster than male mice starting in the 3rd stage of the experiment. (**D**) Average speeds by APP mice split by sex. (**E**) Average speeds by male animals split by groups. Male APP mice run significantly faster than male nTG mice. (**F**) Average speeds by female animals split by groups. Female APP mice run significantly faster than female nTG mice early on in the experiment but nTG mice run faster than female APP mice during the last epoch (days 53-70). Data are presented as the daily means (± SEM) across the entire experiment. Vertical dashed lines represent stage transitions.

**Supplementary Figure 4.**
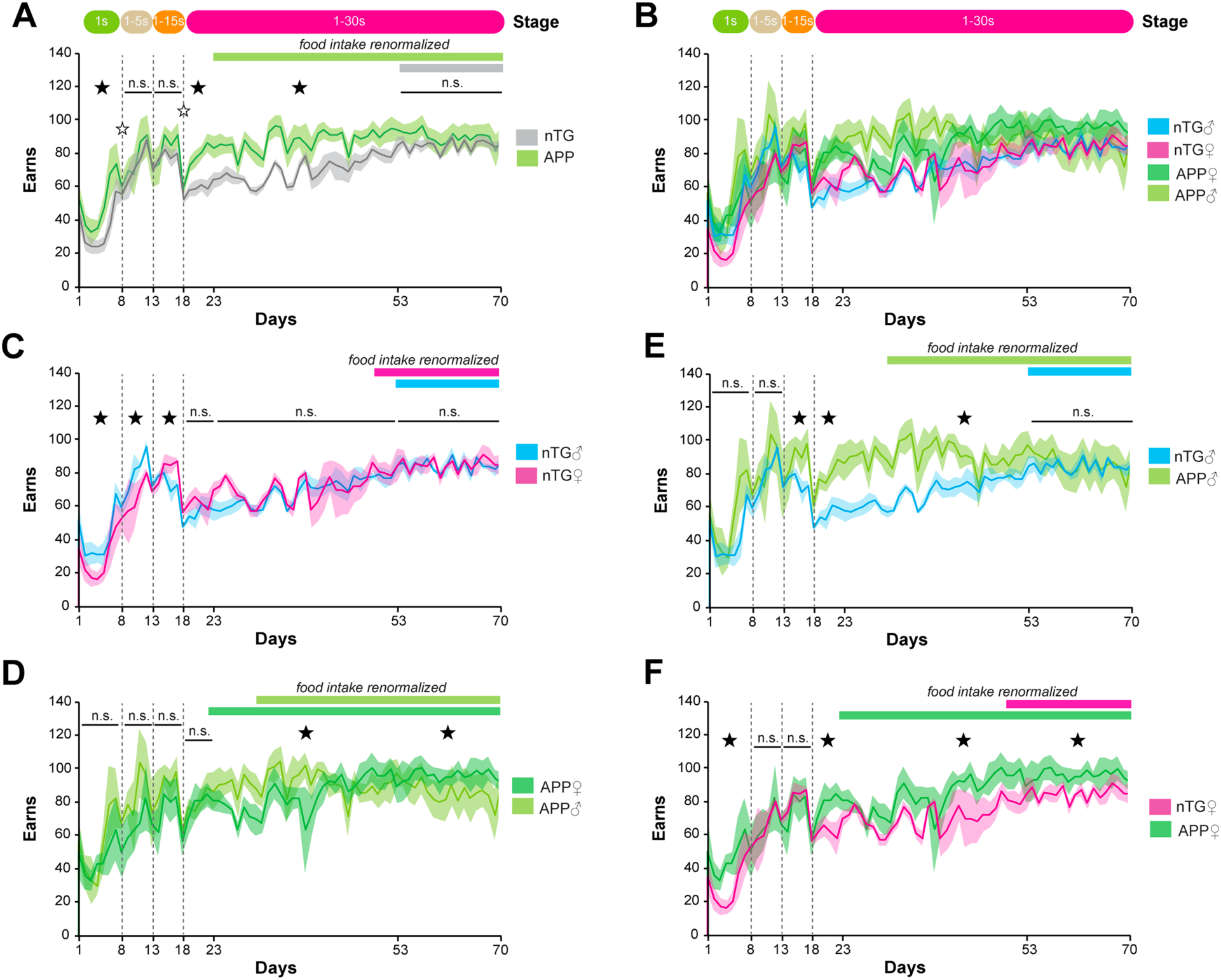
Total amount of pellets earned. (**A**) APP mice earn more pellets than nTG mice. (**B**) Illustration showing earn groups split by sex. (**C**) Male and female nTG mice take turns earning more pellets initially but by the last stage of the experiment male and female nTG mice earn the same amount. (**D**) Male APP mice earn more pellets than female APP mice during the second epoch of the last stage (days 23-52) with female APP mice earning more than male APP mice during the last epoch (days 53-70). (**E**) Male APP mice earn more pellets than male nTG during the first two epochs in the last stage (days 18-22 and days 23-52) but this equalizes at the end of the experiment (days 53-70). (**F**) Female APP mice earn more than female nTG mice. Data are presented as the daily means (± SEM) across the entire experiment. Vertical dashed lines represent stage transitions.

**Supplementary Figure 5.**
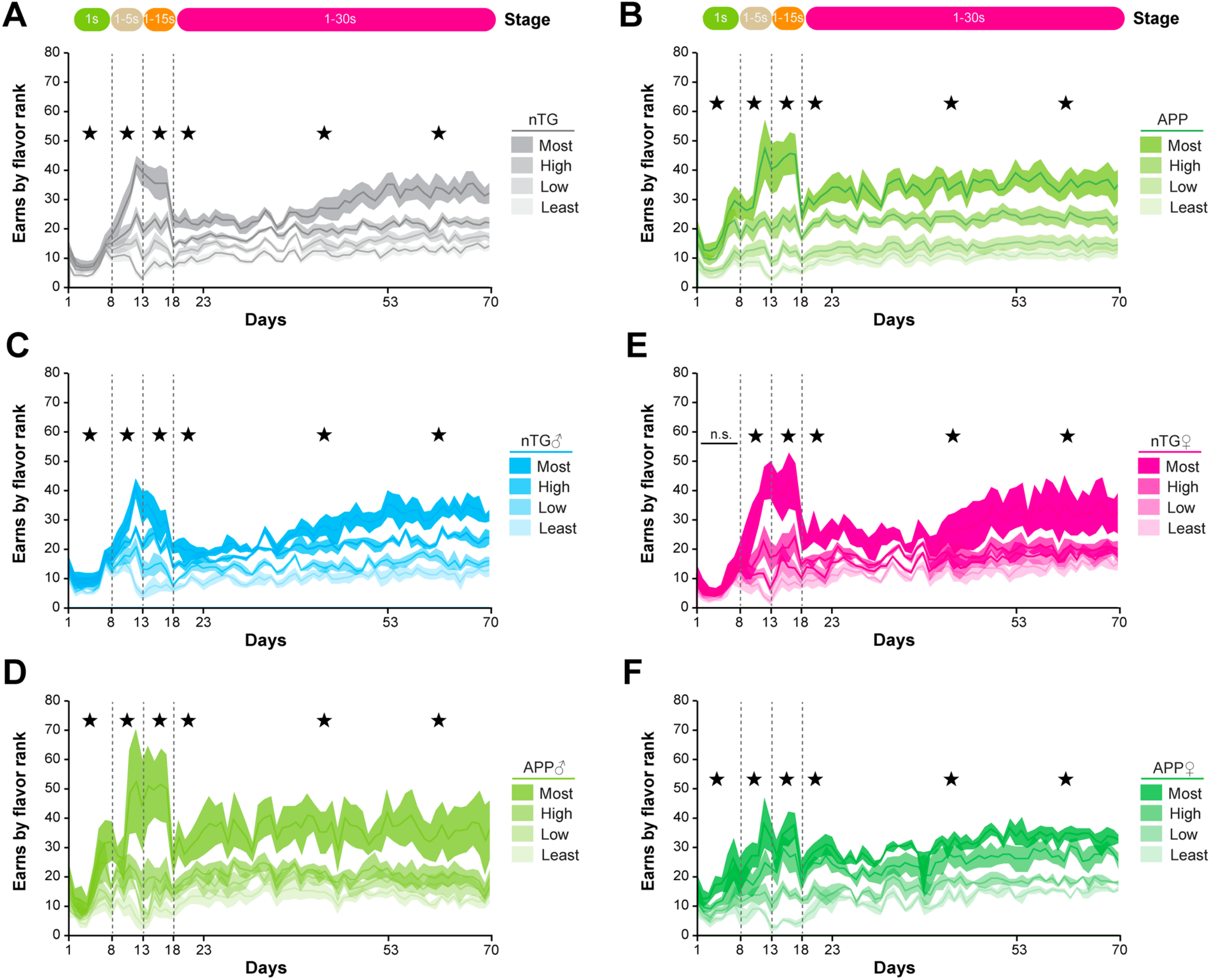
Earns by flavor preferences. To assess flavor preferences, the total earnings of each flavor at the end of the session were examined. Flavors were ranked from most earned to least earned for each individual mouse. (**A**) Earns by rank for nTG mice. (**B**) Earns by rank for APP mice. (**C**) Earns by rank for male nTG mice. (**D**) Earns by rank for male APP mice. (**E**) Earns by rank for female nTG mice. (**F**) Earns by rank for female APP mice. Data are presented as the daily means (± SEM) across the entire experiment. Vertical dashed lines represent stage transitions.

**Supplementary Figure 6.**
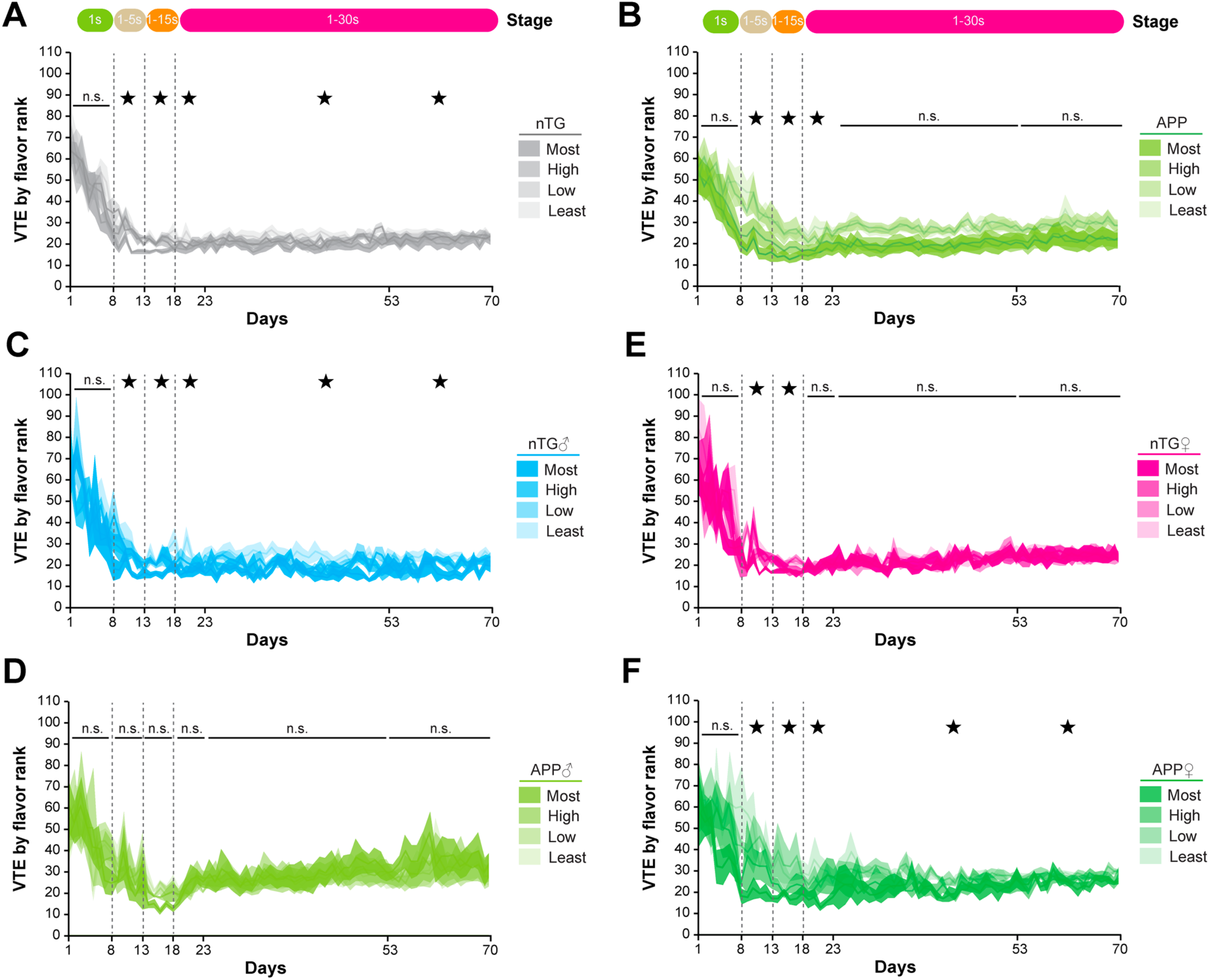
Vicarious-trial-and-error by flavor preferences. (**A**) nTG mice show more VTE for their least preferred flavor and the lowest amount of VTE for their most preferred flavor, as has been reported previously. (**B**) APP mice initially show more VTE for their least preferred and lower VTE for their most preferred flavor. However, they do not show differences in VTE behavior by flavor rank during the last epochs of the last stage (days 23-70). (**C**) Male nTG mice show consistent differences in VTE depending on the flavor. (**D**) Male APP mice do not show distinct levels of VTE for different flavor rankings at any point in the experiment. (**E**) Female nTG mice do not show reliable differences in VTE by flavor rank. (**F**) Female APP mice show distinct levels of VTE for different flavor rankings.

**Supplementary Figure 7.**
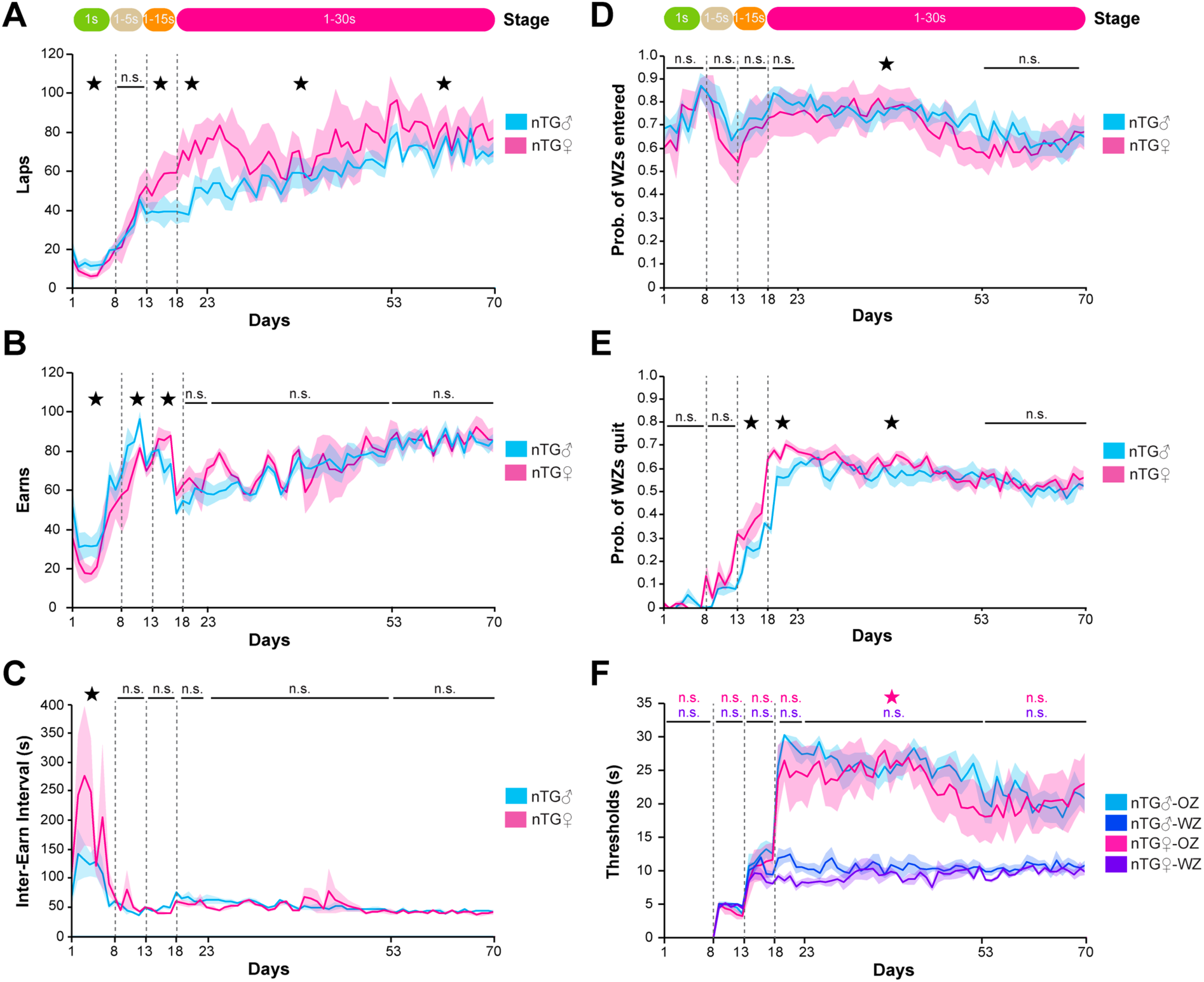
nTG behaviors split by sex. (**A**) Laps run in the correct counterclockwise fashion. Female nTG mice run more laps than male nTG mice. (**B**) Earns per day. (**C**) Inter-earn-interval as defined by the time between earns in seconds. (**D**) Probability of wait zones (WZs) entered. (**E**) Probability of wait zones (WZs) quit. (**F**) Offer zone (OZ) decision thresholds and wait zone (WZ) decision thresholds as a function of cost (offer delay). Star designates p<0.05 for comparisons between groups (male versus female). Data are presented as the daily means (± SEM) across the entire experiment. Vertical dashed lines represent stage transitions.

**Supplementary Figure 8.**
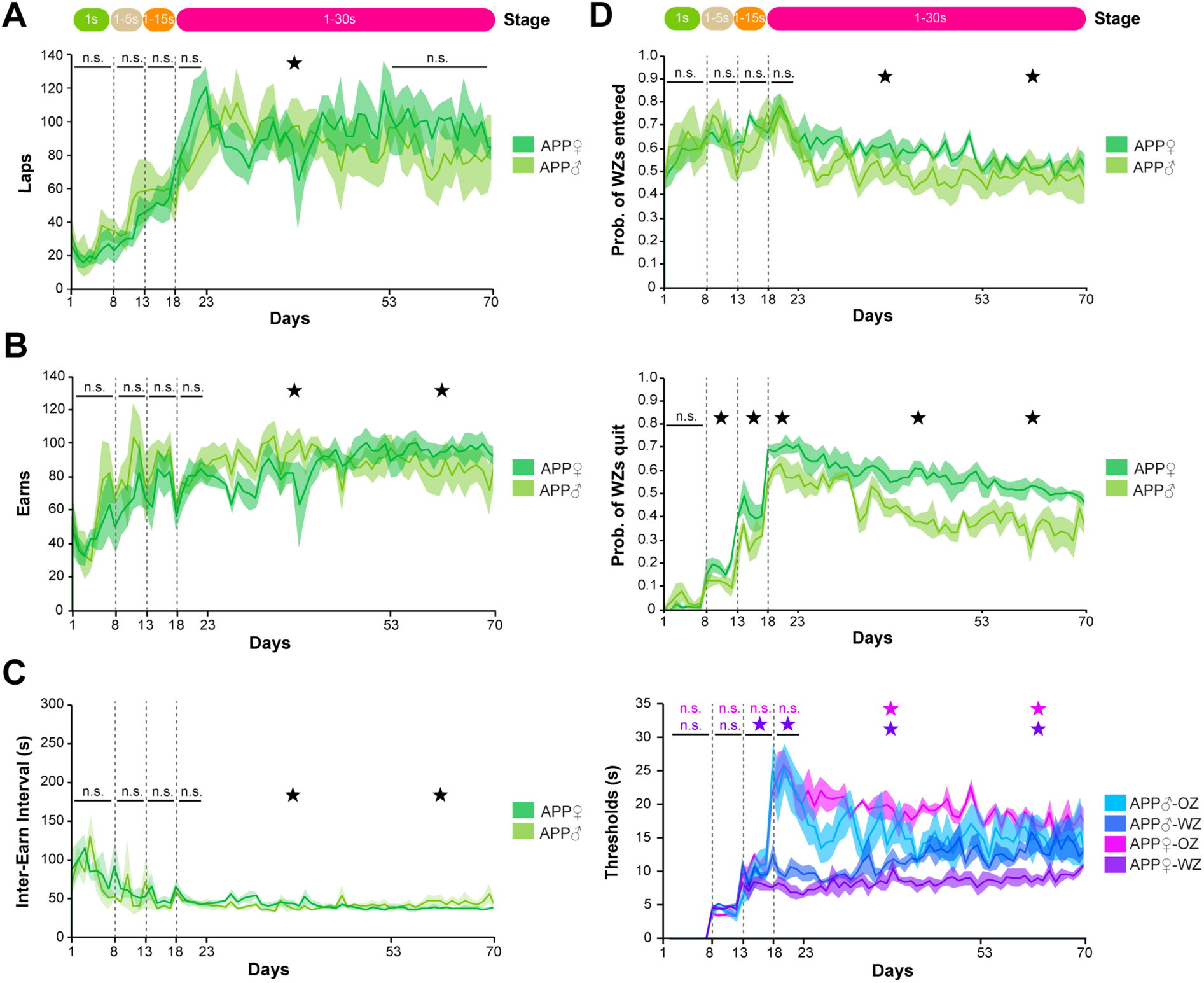
APP behaviors split by sex. (**A**) Laps run in the correct counterclockwise fashion. (**B**) Earns per day. (**C**) Inter-earn-interval as defined by the time between earns in seconds. (**D**) Probability of wait zones (WZs) entered. (**E**) Probability of wait zones (WZs) quit. (**F**) Offer zone (OZ) decision thresholds and wait zone (WZ) decision thresholds as a function of cost (offer delay). Star designates p<0.05 for comparisons between groups (male versus female). Data are presented as the daily means (± SEM) across the entire experiment. Vertical dashed lines represent stage transitions.

**Supplementary Figure 9.**
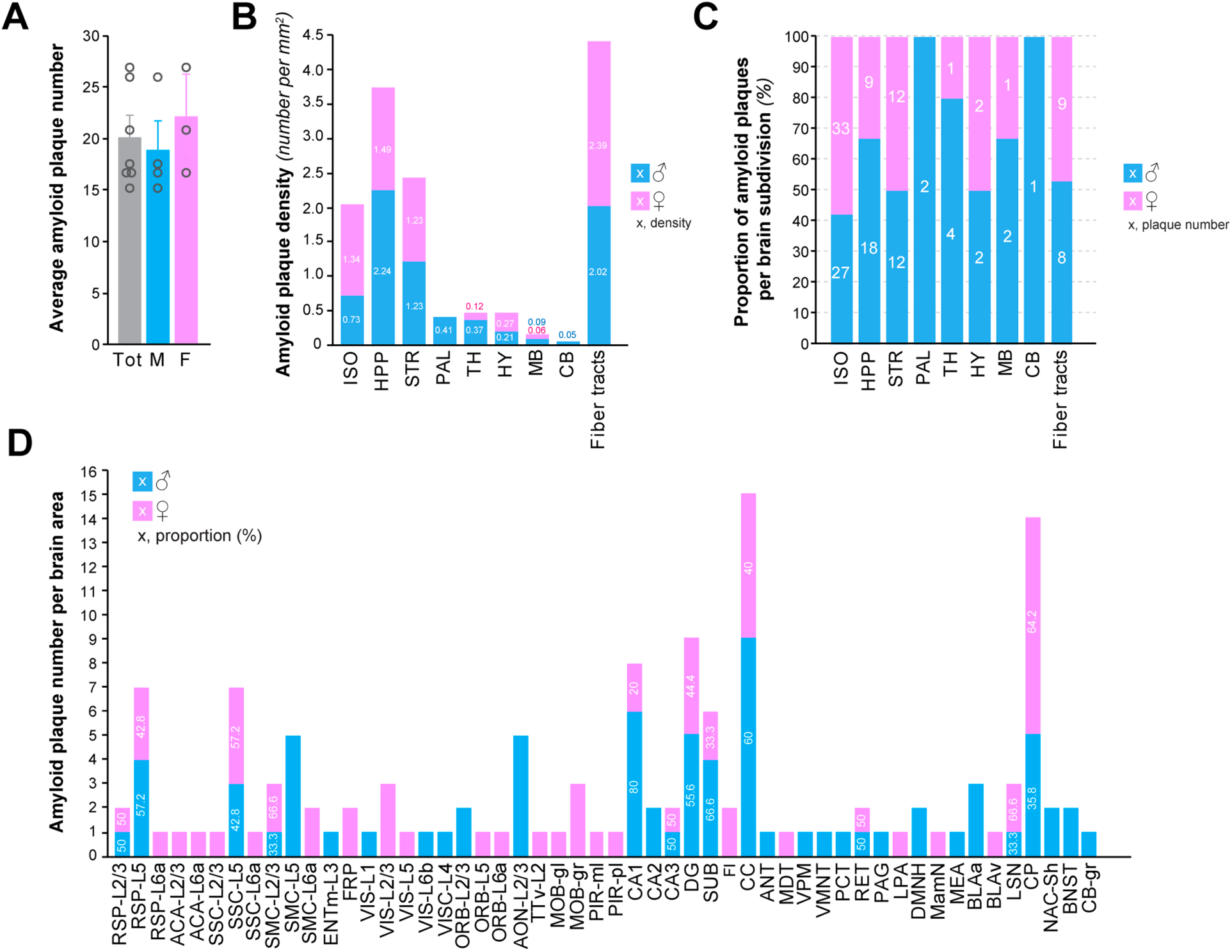
Effect of sex of amyloid pathology in APP mice. (**A**) Quantification of total average plaque numbers. Bars represent mean ± SEM; n = 7 APP mice, including 4 male and 3 female animals. (**B**) Aβ plaque density across brain subdivisions (isocortex [ISO], hippocampus [HPP], striatum [STR], pallidum [PAL], thalamus [TH], hypothalamus [HY], midbrain [MB], cerebellum [CB] and fiber tracts). (**C**) Proportion of amyloid plaques per brain subdivision. (**D**) Amyloid plaque number by brain area, referenced to the Allen Brain Atlas (Lein et al., 2007). Blue bars correspond to male animals while pink bars represent female mice.

**Supplementary Figure 10.**
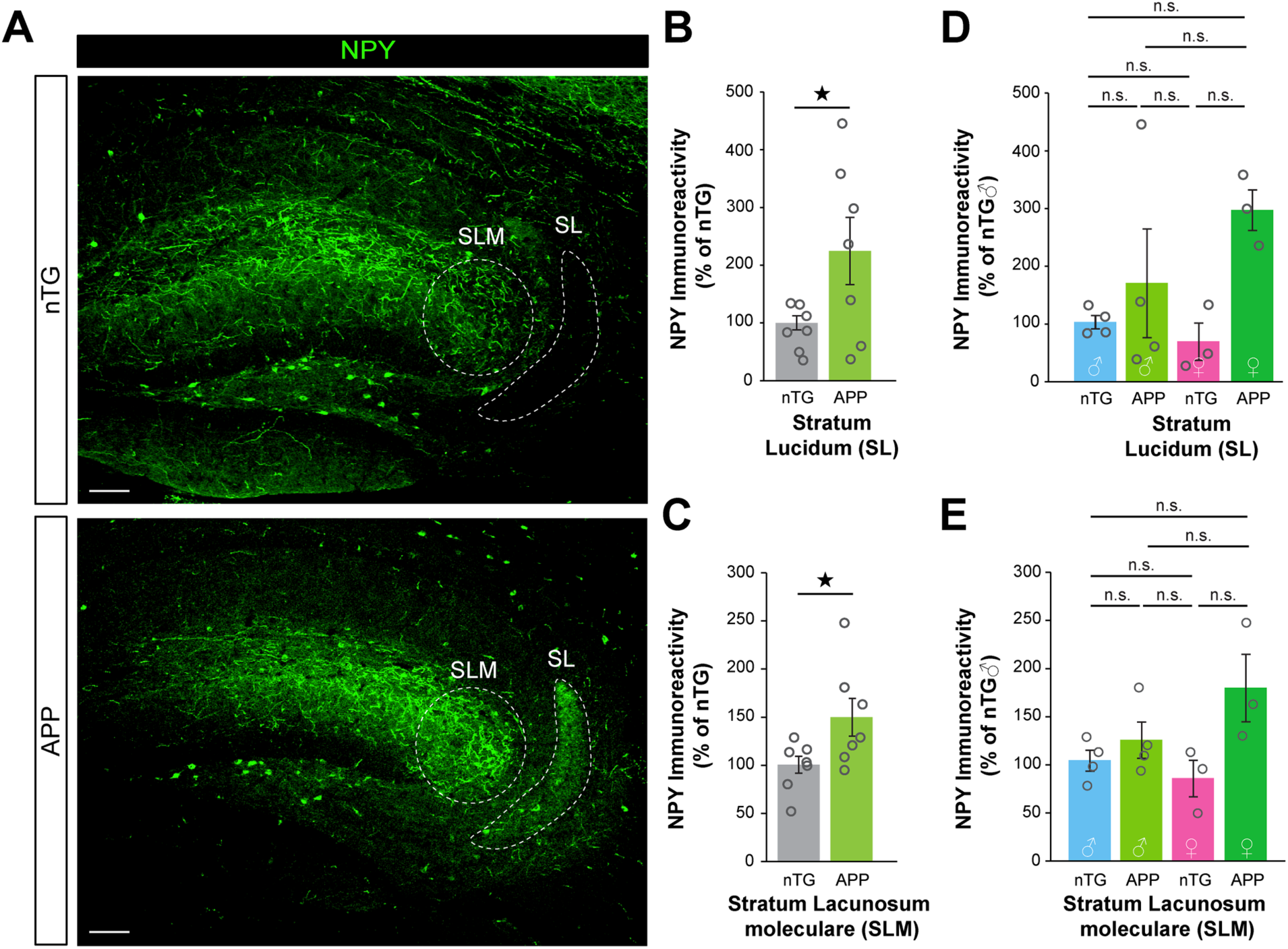
Ectopic expression of NPY in the hippocampi of APP mice. (**A**) Representative immunostaining of the hippocampus from nTG and APP animals stained for neuropeptide-y (NPY, green). (**B,C**) Quantification of NPY immunoreactivity in the (*B*) stratum lucidum (SL) and (*C*) stratum lacunosum moleculare (SLM) of nTG and APP mice (nTG n = 8, APP n = 7, SL *t* test, F_(1,15)_ = 3.5321, P = 0.0414; SLM *t* test, F_(1,15)_ = 4.4108, P = 0.0279). (**D, E**) Quantification of NPY immunoreactivity in the (*D*) SL and (*E*) SLM in the same cohort with groups divide by sex (nTG male n = 4, APP male n = 4, nTG female n = 3, APP female n = 3; SL two-way ANOVA, F_(3,15)_ = 1.8615, P = 0.1945; SLM two-way ANOVA, F_(3,15)_ = 2.6094, P = 0.1041). Data are presented as mean ± SEM.

## REFERENCES

1. Dubois, B. et al. Preclinical Alzheimer’s disease: Definition, natural history, and diagnostic criteria. Alzheimers Dement 12, 292–323 (2016).

2. Coughlan, G., Laczó, J., Hort, J., Minihane, A.-M. & Hornberger, M. Spatial navigation deficits - overlooked cognitive marker for preclinical Alzheimer disease? Nat Rev Neurol 14, 496–506 (2018).

3. Bayard, S., Jacus, J.-P., Raffard, S. & Gély-Nargeot, M.-C. Conscious Knowledge and Decision Making Under Ambiguity in Mild Cognitive Impairment and Alzheimer Disease. Alzheimer Dis Assoc Disord 29, 357–359 (2015).

4. Davis, R., Ziomkowski, M. K. & Veltkamp, A. Everyday Decision Making in Individuals with Early-Stage Alzheimer’s Disease: An Integrative Review of the Literature. Res Gerontol Nurs 10, 240–247 (2017).

5. Han, S. D., Boyle, P. A., James, B. D., Yu, L. & Bennett, D. A. Mild cognitive impairment is associated with poorer decision-making in community-based older persons. J Am Geriatr Soc 63, 676–683 (2015).

6. Denburg, N. L., Tranel, D. & Bechara, A. The ability to decide advantageously declines prematurely in some normal older persons. Neuropsychologia 43, 1099–1106 (2005).

7. Wood, S., Busemeyer, J., Koling, A., Cox, C. R. & Davis, H. Older adults as adaptive decision makers: evidence from the Iowa Gambling Task. Psychol Aging 20, 220–225 (2005).

8. Bennett, D. A. et al. Religious Orders Study and Rush Memory and Aging Project. J. Alzheimers Dis. 64, S161–S189 (2018).

9. Gleichgerrcht, E., Ibáñez, A., Roca, M., Torralva, T. & Manes, F. Decision-making cognition in neurodegenerative diseases. Nature Reviews Neurology 6, 611–623 (2010).

10. Herz, D. M., Bogacz, R. & Brown, P. Neuroscience: Impaired Decision-Making in Parkinson’s Disease. Curr. Biol. 26, R671–673 (2016).

11. Manes, F. F. et al. Frontotemporal dementia presenting as pathological gambling. Nat Rev Neurol 6, 347–352 (2010).

12. Sinz, H., Zamarian, L., Benke, T., Wenning, G. K. & Delazer, M. Impact of ambiguity and risk on decision making in mild Alzheimer’s disease. Neuropsychologia 46, 2043–2055 (2008).

13. Daw, N. D., Niv, Y. & Dayan, P. Uncertainty-based competition between prefrontal and dorsolateral striatal systems for behavioral control. Nature Neuroscience 8, 1704–1711 (2005).

14. Rangel, A., Camerer, C. & Montague, P. R. A framework for studying the neurobiology of value-based decision making. Nat Rev Neurosci 9, 545–556 (2008).

15. Redish, A. D. Beyond the Cognitive Map: From Place Cells to Episodic Memory. (Massachusetts Institute of Technology, 1999).

16. Redish, A. D. The Mind within the Brain: How we make Decisions and How Those Decisions Go Wrong. (Oxfor Univ. Press, 2013).

17. Redish, A. D., Schultheiss, N. W. & Carter, E. C. The Computational Complexity of Valuation and Motivational Forces in Decision-Making Processes. Curr Top Behav Neurosci 27, 313– 33 (2016).

18. Daw, N. D., Gershman, S. J., Seymour, B., Dayan, P. & Dolan, R. J. Model-based influences on humans’ choices and striatal prediction errors. Neuron 69, 1204–15 (2011).

19. Masters, C. L. et al. Alzheimer’s disease. Nat Rev Dis Primers 1, 15056 (2015).

20. Thal, D. R. et al. Alzheimer-related tau-pathology in the perforant path target zone and in the hippocampal stratum oriens and radiatum correlates with onset and degree of dementia. Exp Neurol 163, 98–110 (2000).

21. Villemagne, V. L. et al. Amyloid β deposition, neurodegeneration, and cognitive decline in sporadic Alzheimer’s disease: a prospective cohort study. The Lancet Neurology 12, 357– 367 (2013).

22. Edmonds, E. C. et al. Patterns of Cortical and Subcortical Amyloid Burden across Stages of Preclinical Alzheimer’s Disease. J Int Neuropsychol Soc 22, 978–990 (2016).

23. Hanseeuw, B. J. et al. PET staging of amyloidosis using striatum. Alzheimers Dement 14, 1281–1292 (2018).

24. McDade, E. et al. Cerebral perfusion alterations and cerebral amyloid in autosomal dominant Alzheimer disease. Neurology 83, 710–717 (2014).

25. Sasaguri, H. et al. APP mouse models for Alzheimer’s disease preclinical studies. EMBO J 36, 2473–2487 (2017).

26. Jankowsky, J. L. & Zheng, H. Practical considerations for choosing a mouse model of Alzheimer’s disease. Molecular Neurodegeneration 12, 89 (2017).

27. Mucke, L. et al. High-level neuronal expression of AB1-42 in Wild-Type Human Amyloid Protein Precursor Transgenic Mice: Synaptotoxicity without plaque formation. J Neurosci 20, 4050–4058 (2000).

28. Chin, J. et al. Fyn kinase induces synaptic and cognitive impairments in a transgenic mouse model of Alzheimer’s disease. J. Neurosci. 25, 9694–9703 (2005).

29. Meilandt, W. J. et al. Enkephalin elevations contribute to neuronal and behavioral impairments in a transgenic mouse model of Alzheimer’s disease. J. Neurosci. 28, 5007–5017 (2008).

30. Palop, J. J. et al. Aberrant excitatory neuronal activity and compensatory remodeling of inhibitory hippocampal circuits in mouse models of Alzheimer’s disease. Neuron 55, 697–711 (2007).

31. Cheng, I. H. et al. Accelerating amyloid-beta fibrillization reduces oligomer levels and functional deficits in Alzheimer disease mouse models. J. Biol. Chem. 282, 23818–23828 (2007).

32. Harris, J. A. et al. Transsynaptic progression of amyloid-β-induced neuronal dysfunction within the entorhinal-hippocampal network. Neuron 68, 428–441 (2010).

33. Whitesell, J. D. et al. Whole brain imaging reveals distinct spatial patterns of amyloid beta deposition in three mouse models of Alzheimer’s disease. J Comp Neurol 527, 2122–2145 (2019).

34. van der Meer, M. A., Johnson, A., Schmitzer-Torbert, N. C. & Redish, A. D. Triple dissociation of information processing in dorsal striatum, ventral striatum, and hippocampus on a learned spatial decision task. Neuron 67, 25–32 (2010).

35. Tolman, E. C. Purposive behavior in animals and men. (The Century Co, 1932).

36. Johnson, A. & Crowe, D. A. Revisiting Tolman, his theories and cognitive maps. Cognitive Critique 1, 43–72 (2009).

37. Behrens, T. E. J. et al. What Is a Cognitive Map? Organizing Knowledge for Flexible Behavior. Neuron 100, 490–509 (2018).

38. Aronov, D., Nevers, R. & Tank, D. W. Mapping of a non-spatial dimension by the hippocampal-entorhinal circuit. Nature 543, 719–722 (2017).

39. Dezfouli, A. & Balleine, B. W. Habits, action sequences and reinforcement learning. Eur J Neurosci 35, 1036–51 (2012).

40. LeDoux, J. E. & Daw, N. D. Surviving threats: neural circuit and computational implications of a new taxonomy of defensive behaviour. Nat Rev Neurosci 19, 269–282 (2018).

41. Bandler, R. & Shipley, M. T. Columnar Organization in the Midbrain Periaqueductal Gray: Modules for Emotional Expression? Trends Neurosci 17, 379–389 (1994).

42. Steiner, A. P. & Redish, A. D. Behavioral and neurophysiological correlates of regret in rat decision-making on a neuroeconomic task. Nature Neuroscience 17, 995–1002 (2014).

43. Sweis, B. M., Thomas, M. J. & Redish, A. D. Mice learn to avoid regret. PLoS Biol. 16, e2005853 (2018).

44. Sweis, B. M. et al. Sensitivity to ‘sunk costs’ in mice, rats, and humans. Science 361, 178– 181 (2018).

45. van der Meer, M. A. & Redish, A. D. Theta phase precession in rat ventral striatum links place and reward information. J Neurosci 31, 2843–54 (2011).

46. Bizon, J. L. et al. Spatial reference and working memory across the lifespan of male Fischer 344 rats. Neurobiol Aging 30, 646–55 (2009).

47. Barnes, C. A. & McNaughton, B. L. Neurophysiological comparison of dendritic cable properties in adolescent, middle-aged, and senescent rats. Expimental Aging Research 5, 195–206 (1979).

48. Gallagher, M. & Colombo, P. J. Aging: The cholinergic hypothesis of cognitive decline. Curr Opin Neurobiol 5, 161–168 (1995).

49. Gallagher, M., Nagahara, A. H. & Burwell, R. Cognition and hippocampal systems in aging: Animal models. (Oxford University Press, 1995).

50. Samson, R. D. et al. Age differences in strategy selection and risk preference during risk-based decision making. Behav Neurosci 129, 138–48 (2015).

51. Samson, R. D. & Barnes, C. A. Impact of Aging Brain Circuits on Cognition. Eur J Neurosci 37, 1903–1915 (2013).

52. Sanchez, P. E. et al. Levetiracetam suppresses neuronal network dysfunction and reverses synaptic and cognitive deficits in an Alzheimer’s disease model. Proc. Natl. Acad. Sci. U.S.A. 109, E2895–2903 (2012).

53. Larson, M. E. et al. Selective lowering of synapsins induced by oligomeric α-synuclein exacerbates memory deficits. Proc. Natl. Acad. Sci. U.S.A. 114, E4648–E4657 (2017).

54. Khan, S. S. et al. Bidirectional modulation of Alzheimer phenotype by alpha-synuclein in mice and primary neurons. Acta Neuropathol. 136, 589–605 (2018).

55. Sweis, B. M., Redish, A. D. & Thomas, M. J. Prolonged abstinence from cocaine or morphine disrupts separable valuations during decision conflict. Nat Commun 9, 2521 (2018).

56. Wright, A. L. et al. Neuroinflammation and neuronal loss precede Aβ plaque deposition in the hAPP-J20 mouse model of Alzheimer’s disease. PLoS ONE 8, e59586 (2013).

57. Larson, M. E. et al. Soluble α-synuclein is a novel modulator of Alzheimer’s disease pathophysiology. J. Neurosci. 32, 10253–10266 (2012).

58. Redish, A. D. Vicarious trial and error. Nature Reviews Neuroscience 17, 147–159 (2016).

59. Cox, J. & Witten, I. B. Striatal circuits for reward learning and decision-making. Nat. Rev. Neurosci. 20, 482–494 (2019).

60. Balleine, B. W., Delgado, M. R. & Hikosaka, O. The Role of the Dorsal Striatum in Reward and Decision-Making. J. Neurosci. 27, 8161–8165 (2007).

61. Ferretti, M. T. et al. Sex differences in Alzheimer disease — the gateway to precision medicine. Nat Rev Neurol 14, 457–469 (2018).

62. Lin, K. A. et al. Marked gender differences in progression of mild cognitive impairment over 8 years. Alzheimers Dement (N Y) 1, 103–110 (2015).

63. Hua, X. et al. Sex and age differences in atrophic rates: an ADNI study with N=1368 MRI scans. Neurobiol Aging 31, 1463–1480 (2010).

64. Shansky, R. M. Are hormones a “female problem” for animal research? Science 364, 825– 826 (2019).

65. Palop, J. J. & Mucke, L. Network abnormalities and interneuron dysfunction in Alzheimer disease. Nat. Rev. Neurosci. 17, 777–792 (2016).

66. Palop, J. J., Mucke, L. & Roberson, E. D. Quantifying biomarkers of cognitive dysfunction and neuronal network hyperexcitability in mouse models of Alzheimer’s disease: depletion of calcium-dependent proteins and inhibitory hippocampal remodeling. Methods Mol. Biol. 670, 245–262 (2011).

67. Verret, L. et al. Inhibitory interneuron deficit links altered network activity and cognitive dysfunction in Alzheimer model. Cell 149, 708–721 (2012).

68. Johnson, E. C. B. et al. Behavioral and neural network abnormalities in human APP transgenic mice resemble those of App knock-in mice and are modulated by familial Alzheimer’s disease mutations but not by inhibition of BACE1. Mol Neurodegener 15, 53 (2020).

69. Papale, A. E., Stott, J. J., Powell, N. J., Regier, P. S. & Redish, A. D. Interactions between deliberation and delay-discounting in rats. Cogn Affect Behav Neurosci 12, 513–26 (2012).

70. Steiner, A. P. & Redish, A. D. The road not taken: neural correlates of decision making in orbitofrontal cortex. Front Neurosci 6, 131 (2012).

71. Yin, H. H. & Knowlton, B. J. The role of the basal ganglia in habit formation. Nat Rev Neurosci 7, 464–476 (2006).

72. Johnson, A., van der Meer, M. A. & Redish, A. D. Integrating hippocampus and striatum in decision-making. Curr Opin Neurobiol 17, 692–7 (2007).

73. Chiang, A. C. A. et al. Discrete Pools of Oligomeric Amyloid-β Track with Spatial Learning Deficits in a Mouse Model of Alzheimer Amyloidosis. Am J Pathol 188, 739–756 (2018).

